# Tasting from memory: reconstructing taste from visual cues in the human brain

**DOI:** 10.64898/2026.10.08.757687

**Authors:** Shai Berman, Amir Lawen, Sharon Xu, Itamar Kahn, Daphna Shohamy

## Abstract

Taste guides us toward nourishment and away from harm. Yet when we decide what to eat, taste only exists in memory, evoked by a cue. How the brain retrieves taste memories is almost entirely unknown. If memory re-creates experience, it should reproduce the brain’s activation while tasting, even in the absence of taste. We scanned participants while they formed and then retrieved associations between tastes and visual cues. The gustatory cortex was engaged during retrieval and distinguished among remembered tastes, but not using the patterns it had formed while tasting. Patterns persisted instead in lateral parietal cortex, distinguishing individual tastes, not just taste-categories. Activity in the amygdala covaried with this same parietal region and amygdala activity predicted which associations were retained. Taste memory is not simply a re-creation of sensory experience, but a distinct representation, outside the taste system, yet specific enough to guide what we choose to eat.

## Introduction

Taste memory presents a paradox. It is the memory we feel least: we can picture a loved one’s face or hum a familiar melody, but summoning the sourness of a lemon on demand is surprisingly hard. Yet it is the memory we depend on most. A taste, unlike a face or a melody, is never available to inspect before we commit to it; when we decide what to eat, the flavor is not yet in the mouth, and all we have to guide us is what we remember. Choices that determine nutrition, safety, and survival therefore rest on a memory that is barely accessible to introspection. Almost nothing is known about what the brain retrieves when a taste is remembered.

If a taste memory is stored in a specific place, the gustatory cortex is a natural candidate. This region, encompassing middle-anterior dorsal insula and frontal operculum^1–3^, responds to different kinds of tastes and also represents taste information even in the absence of any gustatory input^4–6^. Activity in the gustatory cortex is shaped by learning and directly participates in the formation of cue-taste associations^7–9^. A parallel literature implicates the amygdala in learning the behavioral significance of these associations, largely through its interactions with gustatory cortex^10–14^. Together these findings describe a circuit capable of acquiring cue–taste associations and reactivating them later, and they motivate a straightforward prediction: recalling a taste should reinstate, in the gustatory cortex, the pattern of activity evoked when that taste was in the mouth.

Work on memory in other modalities complicates this prediction. Early human imaging work found that retrieving a memory reactivates sensory regions engaged during perception^15–20^. These findings were extended by approaches asking not simply where activation occurred, but whether the pattern of activity during perception was recapitulated during memory retrieval^21^. Asking that question directly revealed that reactivation and reinstatement are not the same thing: alongside sensory reactivation, memory-evoked representations were found, in abstract form, in lateral parietal cortex^21–27^. On this account a memory is not a faint copy of an experience but a transformed version of it, and the region that holds it is not the region that first registered it.

The prediction that retrieval of taste memories would manifest similarly to other modalities is further complicated by how well we can recall the experience of taste. While people recognize a flavor instantly, few can conjure one at will in the way they can summon a face or a melody; taste and smell are consistently among the weakest modalities on measures of voluntary imagery^28,29^. The introspective sense that a remembered taste is not much like a tasted one may be misleading. But it may also suggest that what is recovered when a taste is recalled is not a sensory representation at all. Whether the transformed representation held outside of sensory cortex is a general property of memory or a feature of the visual system itself is not known.

How a taste is represented in memory, and therefore where in the brain it can be found, has been difficult to establish, largely for methodological reasons. A cue must be dissociable from the taste it predicts, or it will be impossible to disentangle the brain’s representation of the cue rather than the memory. A memory that has lost detail looks much like a memory that has faded. And because a transformed representation may lie anywhere, the search cannot be confined to gustatory regions in advance. These constraints are difficult to satisfy in animal models, where cue–taste associations of this complexity require extended training, valence is hard to separate from sensory identity, and whole-brain coverage is rarely available. Human fMRI, paired with a task designed to satisfy them, makes the question tractable.

We scanned participants while they tasted six taste solutions spanning sweet, sour and bitter, each paired with two visual cues, and again while they retrieved the taste that each cue predicted from the image alone (**Fig. 1**). Pairing each taste with two cues separated the memory from the object that evoked it. A control condition, in which the same images appeared but attention was directed to their visual features, separated remembering from looking. Grouping the tastes into perceptual categories allowed a classifier trained on one set of tastes to be tested on tastes it had never encountered, isolating what participants retained about whether a taste was, for example, sweet or sour from what they retained about which specific taste it was.

**Figure 1.**
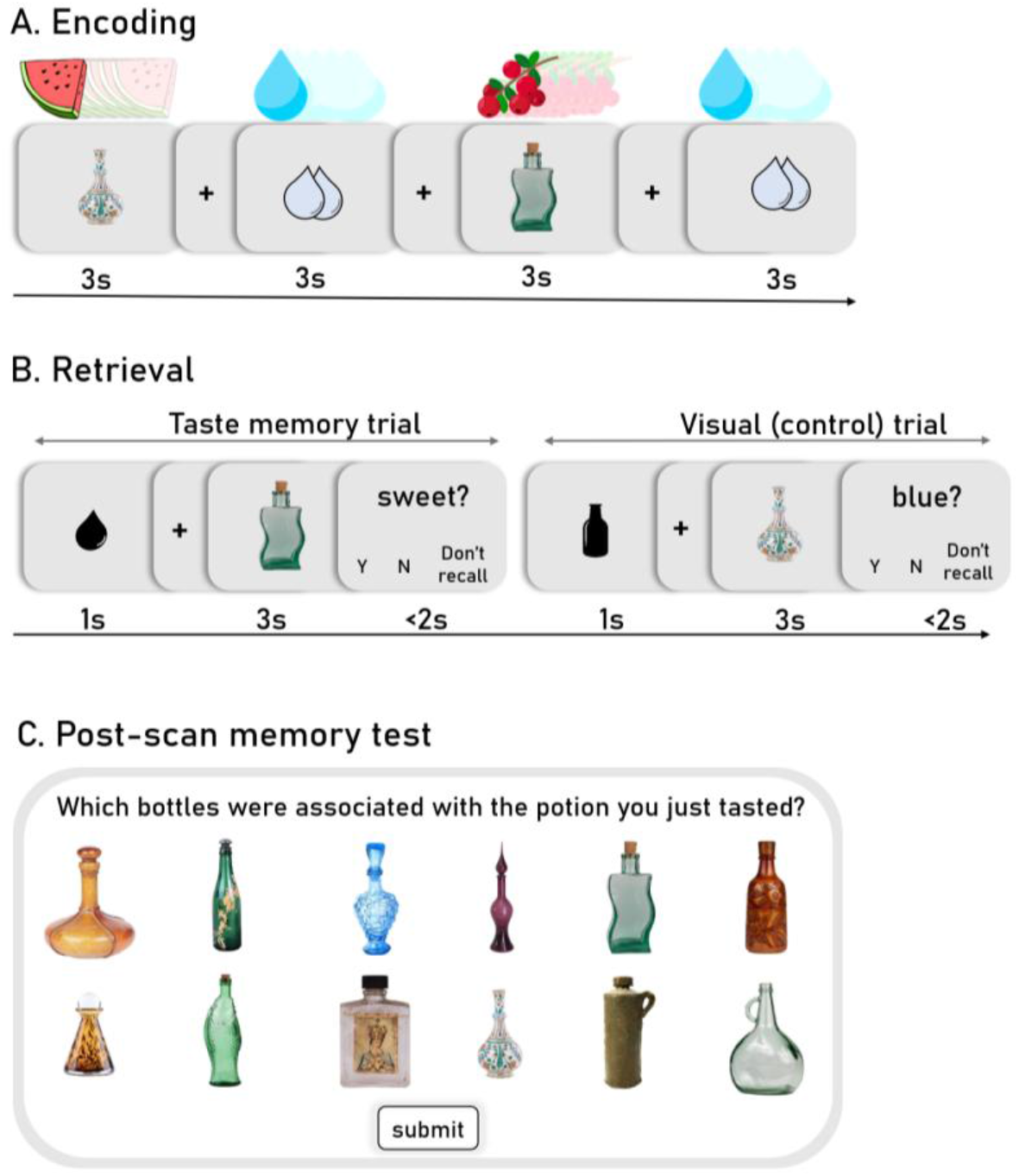
Learning and retrieving cue-taste associations. Participants learned associations between visually distinct bottles and taste solutions, and later retrieved the associated taste from the bottle image alone. Six tastes were used (apple, watermelon, lemon, cranberry, grapefruit, black tea), each paired with two different bottles, giving twelve cue-taste pairs. Pairing each taste with two cues allows taste-related activity to be distinguished from activity driven by any individual image. **(A) Encoding.** A bottle appeared on the screen for 3.5 s with taste delivery (∼0.5 ml) beginning 0.5 s after onset. Each taste-bottle pair was followed by fixation and then by delivery of water accompanied by a rinse cue, which served as the comparison condition for taste delivery: rinse trials matched taste trials in fluid volume, oral stimulation and swallowing, isolating the effect of the taste itself. Encoding included 72 trials across 2 blocks of 8.08 m each. Each bottle-taste pair was presented 6 times, so each taste was delivered 12 times. **(B) Retrieval**. No taste was delivered in this phase. Each trial opened with a 1 s cue indicating its type: a droplet for taste retrieval trials, a bottle icon for visual control trials. A specific bottle then appeared for 3 s, followed by a question about either the taste associated with the bottle or the visual properties of the bottle itself. Participants had 2 s to respond “yes”, “no”, or “don’t recall”. Because both trial types presented the same bottles and required the same responses, differing only in what participants were asked to judge, the control condition isolates retrieval of the association from the perception of a familiar image. This phase consisted of 108 trials across 3 blocks of 8.37 m each (90 taste retrieval trials, 18 visual control). **(C) Post-scan memory test.** Participants rated each taste on sweetness, sourness, bitterness, and pleasantness, and then selected, for each taste, the two bottles it had been paired with from all twelve alternatives.

Training classifiers on activity during tasting and testing them during recall then allowed the central question to be asked directly: is the representation recovered when a taste is remembered the one formed when it was experienced? We found that the gustatory cortex was engaged during retrieval and its activity distinguished among remembered tastes. But, patterns established during tasting in the gustatory cortex did not generalize to retrieval. Instead, we found such generalization in lateral parietal cortex, a region identified in advance from the visual memory literature. Activity in the amygdala during learning predicted which associations were most reliably retained, scaling with how strongly pleasantness distinguished them, and at retrieval amygdala activity covaried with these same parietal regions. These results show that a remembered taste is not a faint version of a tasted one. It is a different representation, in a different place.

## Results

Participants (n=37) learned associations between twelve visual cues and six tastes while undergoing fMRI, and were scanned again while recalling what each cue predicted. Sample sizes vary across analyses below due to exclusions and are given in the figure captions and Methods.

Memory was reliable but not at ceiling. During scanning, participants answered questions about the taste associated with each bottle at 83.2±9.5% accuracy (t(34)=20.77, p<.001). In a post-scan test, they selected the two visual cues paired with each taste from twelve alternatives, succeeding on 57.9±20.3% of trials (t(35)=12.17, p<.001; chance=16.7%). The tastes were selected to span sweet, sour and bitter; as described below, only the sweet and sour groups cohered in participants’ perception and analyses of taste category were restricted accordingly.

### Gustatory cortex is engaged when a taste is recalled, not only when it is tasted

We first asked whether regions that respond to taste are also recruited when a taste is retrieved (**Fig. 2**). During encoding, we contrasted taste delivery with rinse delivery. Taste increased BOLD activity in bilateral gustatory cortex, consistent with previous studies on taste perception^30–32^. Taste also increased activation across a distributed set of regions including visual cortex and subcortical nuclei (**Fig. 2A**; all clusters listed in Table S1). Because rinse and taste delivery were matched in volume, oral stimulation and swallowing, the contrast isolates what distinguishes a taste from rinse not only its gustatory properties but also its palatability, salience, and acquired associations.

**Figure 2.**
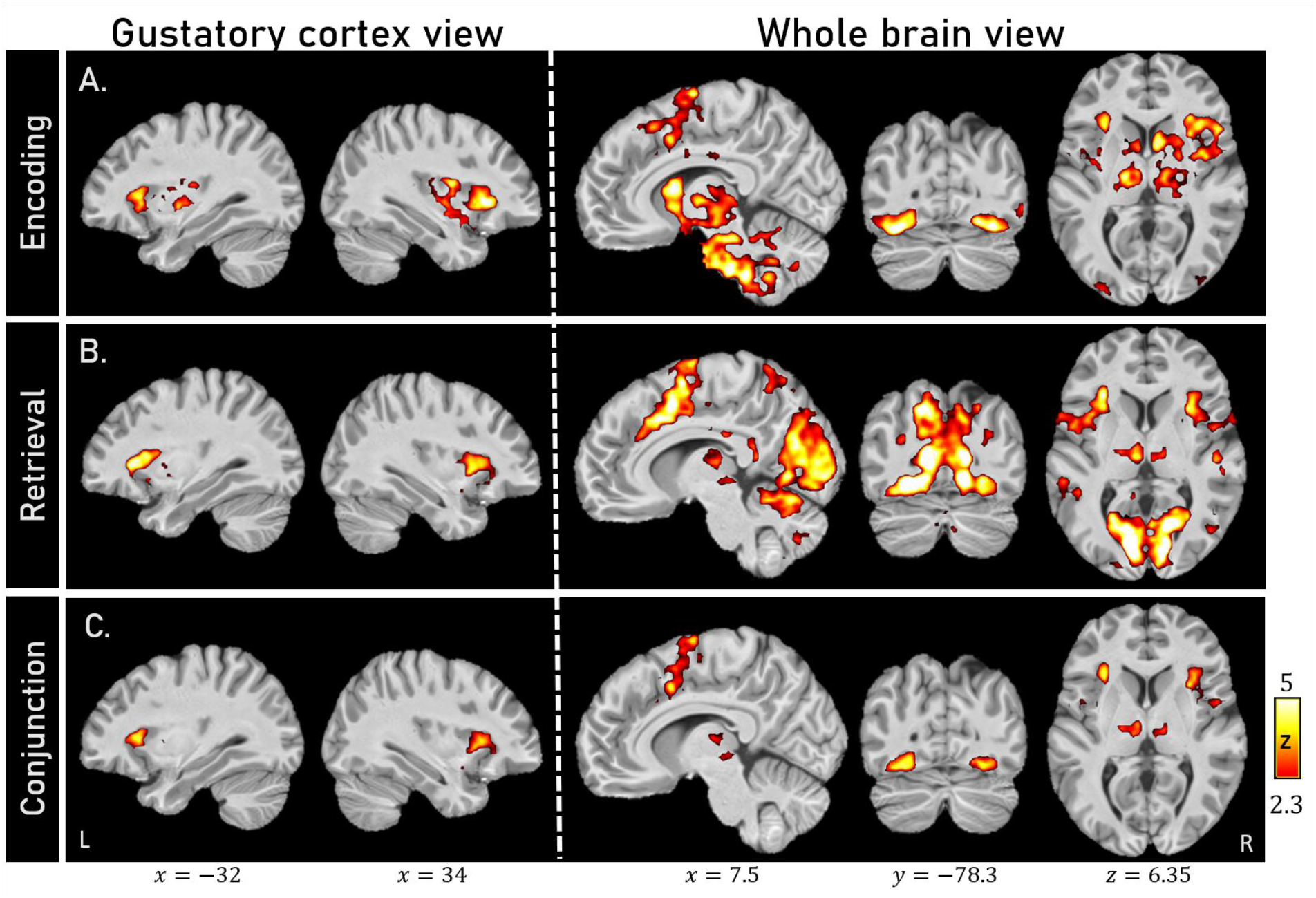
Gustatory cortex is engaged during retrieval as well as during tasting. **(A) Encoding.** Taste delivery contrasted with rinse. Because rinse trials matched taste trials in fluid volume, oral stimulation and swallowing, this contrast isolates the effect of the taste itself. Activation encompassed bilateral gustatory cortex, visual cortex, medial frontal cortex, and extensive subcortical regions including thalamus, caudate, midbrain and brainstem. **(B) Retrieval.** Taste retrieval trials contrasted with visual control trials, estimated over the period in which the bottle was displayed. No taste was delivered in this phase. Both trial types presented the same bottles, differing only in whether participants judged the associated taste or the image itself. Activation included gustatory cortex, visual cortex, medial frontal cortex, thalamus, and cerebellum. **(C) Conjunction of encoding and retrieval**. The conjunction was computed as the voxel-wise minimum of the two maps. Overlap was limited to gustatory cortex, visual cortex, thalamus, and medial frontal cortex.

During retrieval, no taste was delivered: participants saw a cue and were asked about the characteristics of the taste associated with it (e.g., its sweetness or fruitiness). Activity in bilateral gustatory cortex nonetheless increased relative to a control condition in which the same cues appeared but participants judged their visual properties (**Fig. 2B**). Retrieval also engaged the visual cortex, medial frontal cortex, thalamus and cerebellum. This effect does not reflect visual input or motor demand, since both conditions presented the same images and required the same responses. Instead, it reflects what participants attended to: the associated taste in one case, the image itself in the other.

The two contrasts produced distributed activation, but their overlap was limited. A conjunction analysis identified gustatory cortex, lateral occipital cortex, thalamus, and medial frontal cortex (**Fig. 2C**).

The left images show sagittal views of masked brain activation to facilitate identification of gustatory cortex activation. The right maps show whole-brain activation maps cluster-corrected at z > 2.3. n=36 for all three analyses. Full cluster listings, including coordinates and extents, are given in Table S1.

### Taste identity is represented during retrieval in a different network than during tasting

Activation magnitude alone cannot separate regions representing taste from ones that are more generally engaged in the task. To examine whether activity patterns in these regions distinguish among tastes, we trained a linear classifier to distinguish each taste from the others, with cue identity controlled so that visual information could not support classification.

During tasting, taste identity was decodable from patterns in frontal regions, including orbitofrontal cortex (OFC) and ventrolateral prefrontal cortex (PFC), and in sensory and sensorimotor areas, including peri-central cortex and a cluster in the gustatory cortex (**Fig. 3A**). During retrieval, when no taste was delivered, taste identity was again decodable—but from a partly different set of regions, including right gustatory cortex and association and cingulate cortex (**Fig. 3B**).

**Figure 3.**
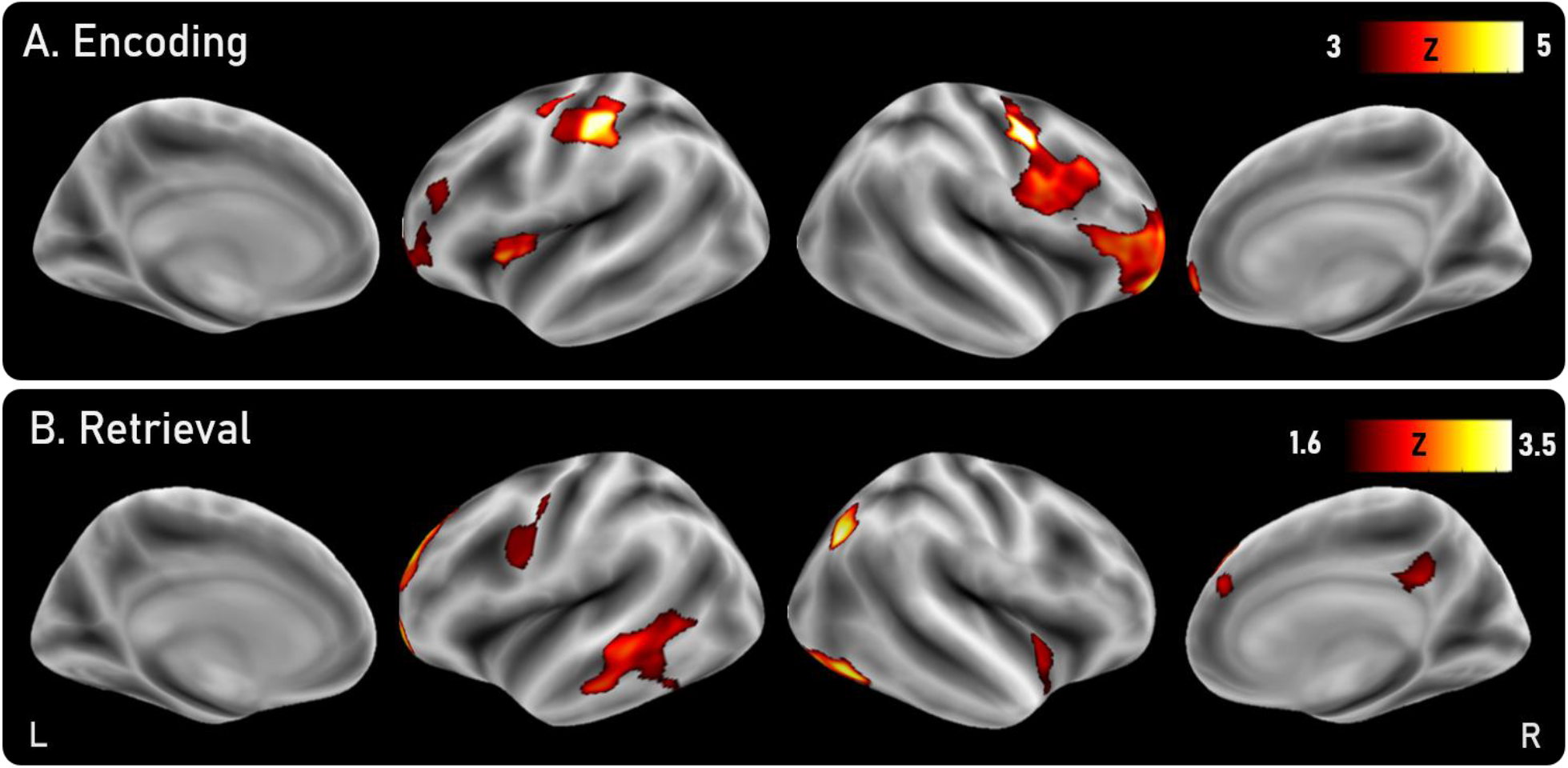
Taste identity is decodable during both encoding and retrieval. **(A) Encoding**. Decoding accuracy was above chance in the gustatory cortex, OFC, ventrolateral PFC, peri-central cortex, primary sensory and premotor cortex. **(B) Retrieval**. Decoding accuracy was above chance in the right lateral parietal cortex, right gustatory cortex, right cingulate cortex and in right temporo-occipital cortex, as well as in left pre-central cortex, left superior-frontal regions, and left temporal cortex. Results are whole brain maps of taste identity decoding computed with a surface-based searchlight, projected on the fsaverage6 surface; color indicates z-statistics from the permutation test against chance (1/6); maps were cluster-corrected using permutation-based cluster-level familywise error correction. For encoding (n=32), where the effects were strong, we used a voxel threshold of p<0.001, and for retrieval (n=28) p<0.05. Maps are smoothed for visualization (statistics computed on unsmoothed data). Full cluster listings are given in Table S2. Encoding accuracy maps with a voxel threshold of p<0.05 are presented in supplementary Figure S1.

Taste-specific information was therefore present during memory retrieval, in the absence of any gustatory input: what participants recovered from a cue was not merely that it had been paired with tasting, but the specific taste it was paired with. The regions carrying that information, however, are not the same as those carrying it during tasting. Association and cingulate cortex were informative during retrieval, but not during tasting, while much of the sensorimotor network was only informative during tasting, but not during retrieval.

### The code that bridges tasting and remembering lies outside the gustatory cortex

A region can be informative when a taste is in the mouth, and again later when it is being remembered, but may do so using unrelated codes in each. Neither the overlap between phases nor its absence can be interpreted without testing whether a pattern learned in one phase generalizes to the other.

We therefore trained classifiers on encoding activity and tested them on retrieval activity, requiring that patterns learned while a taste was in the mouth generalize to a period in which no taste was delivered. We refer to this as *cross-phase generalization*. Across the cortex, cross-phase generalization was successful in the lateral parietal cortex, as well as in OFC and in the middle temporal gyrus (**Fig. 4A**). Patterns established while a taste was in the mouth were sufficient to classify what participants later recalled from a visual cue alone, but the regions supporting this transfer lay outside of the gustatory regions engaged during tasting.

**Figure 4.**
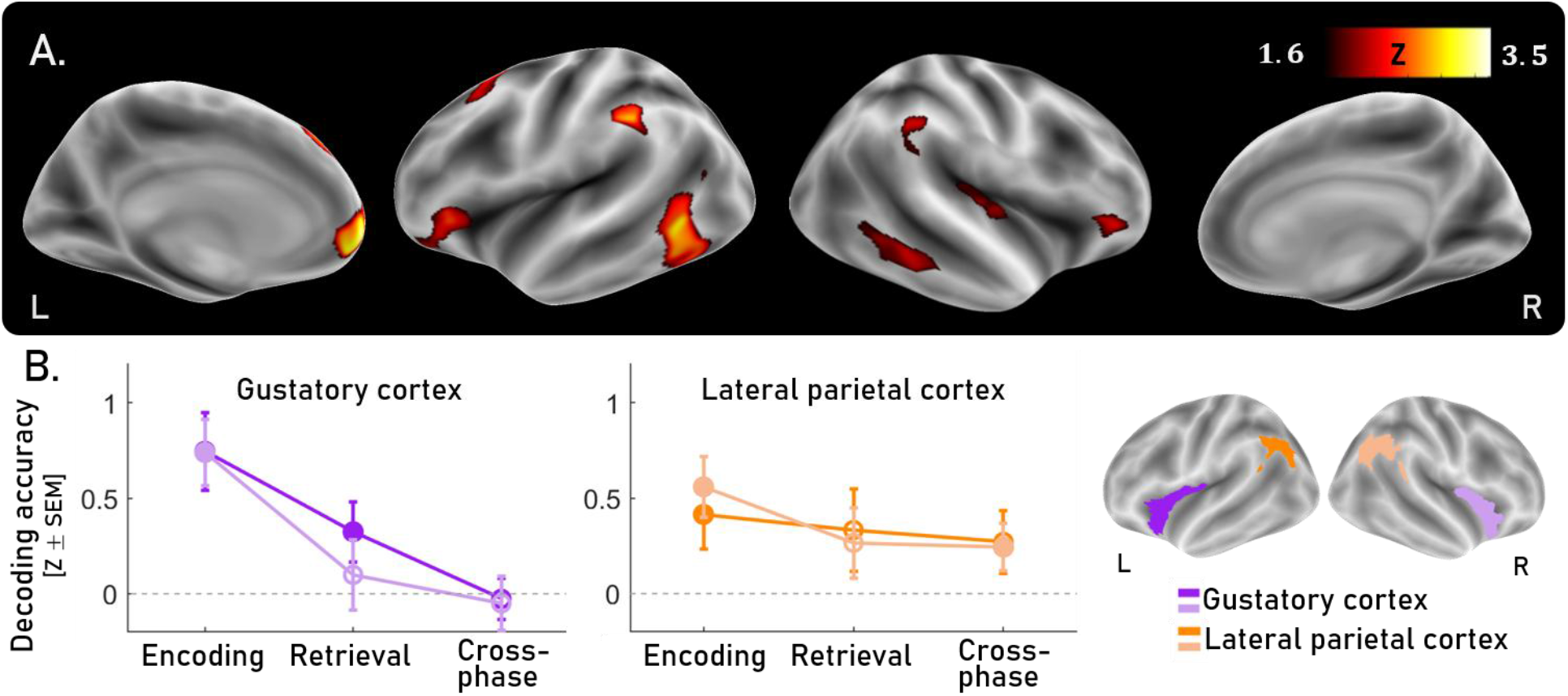
Taste representations generalize from encoding to retrieval. **(A) Taste identity, whole brain.** Cross-phase decoding was above chance in right OFC, medial PFC, and middle temporal gyrus, and lateral parietal cortex, in both hemispheres, as well as in a posterior portion of right insula, and left superior frontal cortex left angular gyrus, visual association cortex, ventral posterior cingulate cortex, left OFC and left inferior frontal cortex. Classifiers were trained on encoding activity and tested on retrieval activity, when no taste was delivered. Category classifiers were trained on one taste from each category and tested on the held-out pair, so decoding cannot reflect individual taste identity. **(B) Decoding taste identity in regions-of-interest.** Decoding taste identity in gustatory cortex and lateral parietal cortex ROIs (specifically, anterior dorsal insula, in purple, and angular gyrus in orange), defined anatomically and specified in advance, shown separately for each hemisphere. The accuracy is plotted for encoding, retrieval and cross-phase decoding. All ROIs represented taste identity during encoding. The left gustatory cortex also showed taste identity representation at retrieval but showed no evidence of cross-phase generalization at any feature count. The lateral parietal cortex did present generalization across learning phases. Filled symbols indicate p < .05; error bars show SEM. The ROIs are displayed on the right, on the fsaverage6 inflated brain.

In addition to this whole brain analysis, we also tested cross-phase generalization in two regions specified in advance: the gustatory cortical area, specifically anterior dorsal insula, where a reinstated sensory code would be expected, and the lateral parietal cortex, specifically the angular gyrus, which the visual memory literature identifies as a site of transformed rather than reinstated retrieval codes^24–26,33^ (**Fig. 4b**). In each ROI, decoding was performed across a range of feature-selection thresholds (k) and summarized as a single integrated measure of decoding strength per subject, tested against chance (see Methods). Accuracy across feature-selection thresholds is presented in Supplementary Fig. S2.

Searchlight maps: color indicates z-statistics from permutation testing against chance, projected on the fsaverage6 surface, cluster-corrected using permutation-based cluster-level familywise error correction, smoothed for visualization; statistics computed on unsmoothed data. ROI analyses: feature selection was performed within each cross-validation fold on training data only, applied identically to both regions. Full cluster listings are given in Table S3. n = 32.

In the gustatory cortical region, taste identity was decodable during tasting (t(31)=5.38, p<0.001). The same voxels yielded no evidence of cross-phase generalization (t(31)=-0.43, p=0.7). In the angular gyrus, on the other hand, taste identity was decodable during tasting (t(31)=3.85, p<0.001), and it also generalized from tasting to retrieval, as reflected in successful cross-phase decoding (t(32)=2.29, p=0.03).

A region × phase interaction confirmed this dissociation: the drop from the encoding phase to cross-phase generalization was larger in gustatory cortex than in angular gyrus (F(31)=6.5, p=0.012). Because the two regions were treated identically (same classifier, same feature selection, and same range of feature counts) and because the gustatory cortical regions supported strong identity decoding during tasting, the absence of transfer there is unlikely to reflect insufficient signal or an unsuitable mask.

Together, the whole-brain and ROI results converge: the lateral parietal cluster identified by an unbiased search across the cortex falls within a predetermined region established in the visual memory literature. Both gustatory cortex and lateral parietal cortex represented taste during tasting, but only lateral parietal cortex generalized across phases. In the transition from tasting to remembering, the brain retains the identity of which taste was experienced, but the identity is represented by an activation that no longer resembles the one the gustatory cortex used while tasting it. This suggests that retrieving a taste is not a matter of re-tasting it, and it involves transformed representations.

### Taste category and taste identity are separable, in behavior and in the brain

The six tastes were selected to span sweet, sour, and bitter, but whether participants perceived them that way is an empirical question. We characterized the perceptual structure of the stimulus set from participants’ own ratings, correlating each taste’s profile of sweetness, sourness and bitterness across participants (**Fig. 5A**). Post-scan similarity ratings clustered watermelon with apple and cranberry with lemon, whereas the two putatively bitter stimuli did not form a comparable group; grapefruit grouped weakly with the sour tastes and black tea with nothing. Category analyses were therefore restricted to sweet and sour (all six tastes were retained in the taste identity analyses reported above).

**Figure 5.**
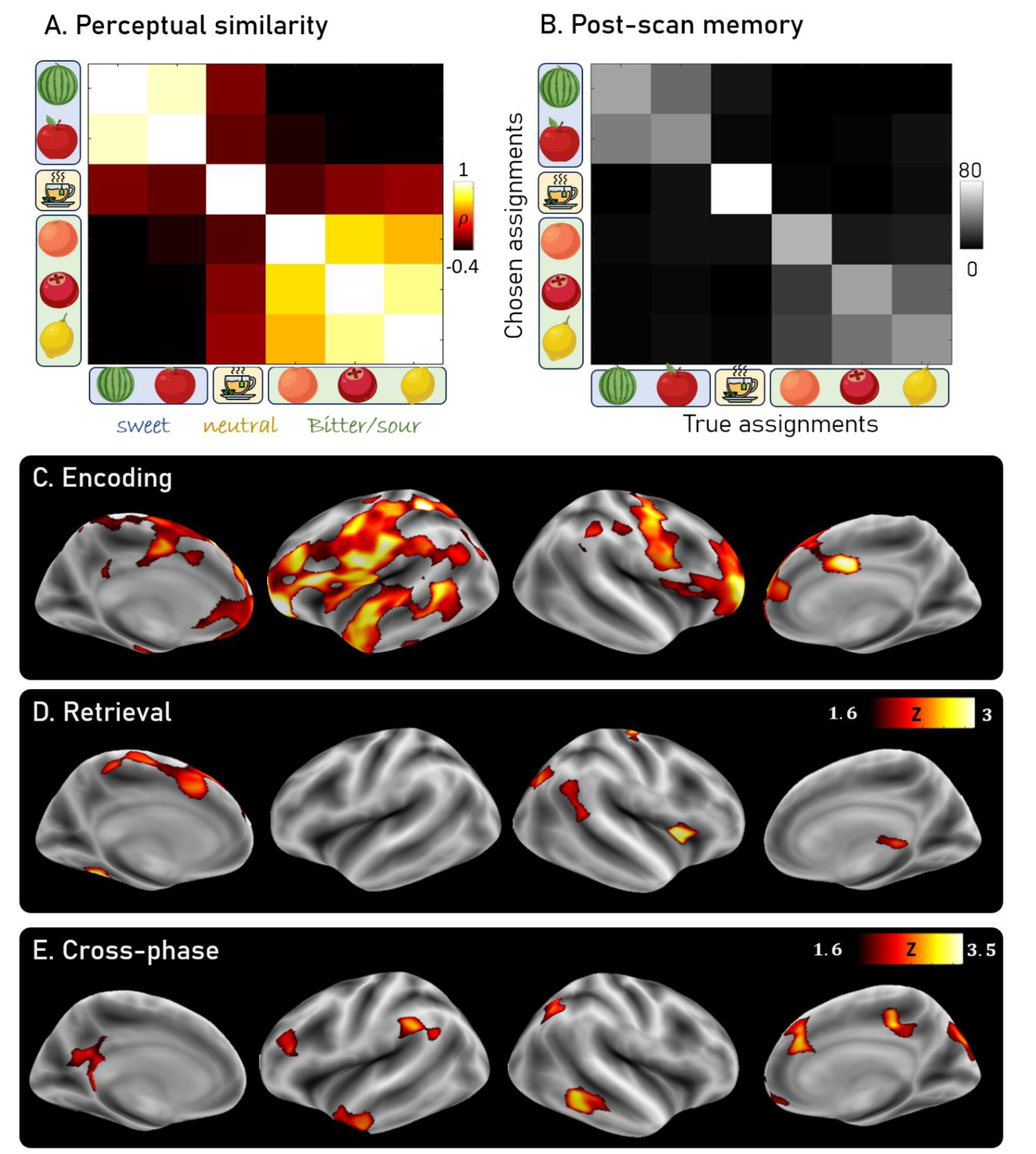
Taste category shapes behavior and is decodable in the brain. **(A) Perceived similarity among tastes.** Similarity was computed as the correlation, across participants, between tastes in their rated sweetness, sourness and bitterness. Two clear groupings emerged: watermelon with apple (sweet) and cranberry with lemon (sour). The two intended bitter tastes did not form a comparable group. (n=37). **(B) Memory errors respect perceptual structure**. Confusion matrix from the post-scan memory test, in which participants selected the two visual cues associated with each taste from twelve alternatives. Errors fell disproportionately on cues paired with a perceptually similar taste. (n=37). **(C) Encoding.** Category was decodable in a widespread set of regions, including gustatory cortex and operculum, peri-central and primary sensory cortex, OFC, lateral and frontopolar PFC, middle anterior cingulate cortex, and left temporal cortex. **(D) Retrieval**. Category was decodable in a more restricted set of regions, including right gustatory cortex, right temporo-parietal and lateral parieto-occipital cluster (both including the angular gyrus), as well as in the left fusiform gyrus, and left medial frontal cortex and anterior cingulate cortex. **(E) Cross-phase.** Classifiers trained on encoding trials and tested on retrieval decoded category above chance bilaterally in angular gyrus and posterior cingulate cortex, right medial frontal cortex, OFC, visual association cortex and middle temporal gyrus, and left temporal pole and dlPFC.

Memory errors in the post-scan test respected the structure of perceived taste categories. Participants who failed to select the correct bottle overwhelmingly chose one paired with a taste from the same perceptual category (**Fig. 5B**). This distinction between memory for the identity vs. the category matters for two reasons. First, taste categories signal broad nutritional properties and a memory that preserves category while losing item identity remains sufficient to guide approach or avoidance. Second, the categorical representation aligns with what a transformed representation may look like behaviorally: intact at a coarse level, degraded at a fine one. We therefore asked whether the categorical structure evident in behavior is present in the brain, and whether it is carried by the same regions that represent taste identity.

We trained a linear classifier to distinguish sweet from sour, training on one taste from each category and testing on the remaining pair. Because the classifier was tested on tastes it had never encountered, successful decoding cannot reflect the identity of any individual taste.

During tasting, taste category was decodable from a widespread network (**Fig. 5C**), with the strongest effects in somatomotor and somatosensory regions including gustatory cortex and peri-central cortex, and in frontal regions including OFC, lateral PFC, and the frontal pole. Category was also decodable in middle anterior cingulate cortex and left temporal pole, where identity decoding did not reach significance.

At retrieval, category decoding did not show a comparable expansion relative to identity decoding and was significant in the right temporo-parietal and parieto-occipital cortex, left fusiform gyrus, right gustatory cortex, and medial frontal cortex (**Fig. 5D**).

Thus, both at encoding and retrieval, patterns in the medial frontal cortex distinguished sweet from sour, but did not discriminate individual tastes, while lateral parietal cortex and right gustatory cortex carried both. Category and identity are therefore separable in the brain, as they are in behavior, and at least one region (medial frontal cortex) represents taste category without representing taste identity.

Applying the cross-phase test to taste-category gave a weaker version of the pattern found for taste-identity. Across the cortex, category generalized from tasting to taste retrieval in a partly overlapping set of regions, including angular gyrus and posterior cingulate cortex, right medial frontal cortex, OFC, visual association cortex and middle temporal cortex (**Fig. 5E**). In the ROI analysis (supplementary Fig. S3), category was decodable during tasting in gustatory cortex (t(31)=5.11, p<0.001), and in lateral parietal cortex (t(31)=2.83, p<0.01). But there was no evidence of cross-phase generalization (gustatory cortex: t(31)=0.8, p=0.4; lateral parietal cortex:(t(31)=0.96, p=0.3). The region × phase interaction was in the same direction as for taste identity, but was not significant (F(32)=2.4 p=0.12).

Imaging results are projected on the fsaverage6 surface; color indicates z-statistics from the permutation test against chance (1/6); maps were corrected using permutation-based cluster-level familywise error correction. For encoding (n=32), where the effects were strong, we used a voxel threshold of p<0.001, and for retrieval (n=31), and cross-phase (n=32), we used a voxel threshold of p<.05. Maps are smoothed for visualization (statistics computed on unsmoothed data). Full cluster listings are given in Table S4. Encoding accuracy maps with a voxel threshold of p<0.05 are presented in supplementary figure S1.

Category information is therefore present in both phases and generalizes between them in the whole-brain analysis, though the evidence that it does so in angular gyrus specifically is weaker than for taste identity: the ROI effect was not significant, and the searchlight clusters fell in nearby lateral parietal regions. In gustatory cortex, by contrast, results were consistent across both levels: neither the identity nor the category code established during tasting was recoverable at retrieval.

### Amygdala activity during encoding predicts which associations are later remembered

The analyses so far describe what taste memories are made of. They do not address why some are retained and others are not. We therefore asked whether activity during encoding predicted which associations participants would later remember, comparing encoding activity for cue–taste pairs subsequently remembered perfectly (100% accuracy) with those for which memory was weaker (<100% accuracy; see Methods).

This subsequent memory effect was significant in the amygdala, where encoding activity was greater for consistently remembered tastes (t(29)=3.18, p<0.01; **Fig. 6A**). No other region survived whole-brain correction. Because the hippocampus is central to the formation of memory associations, we also examined it directly as an ROI, and found no subsequent memory effect there.

**Figure 6.**
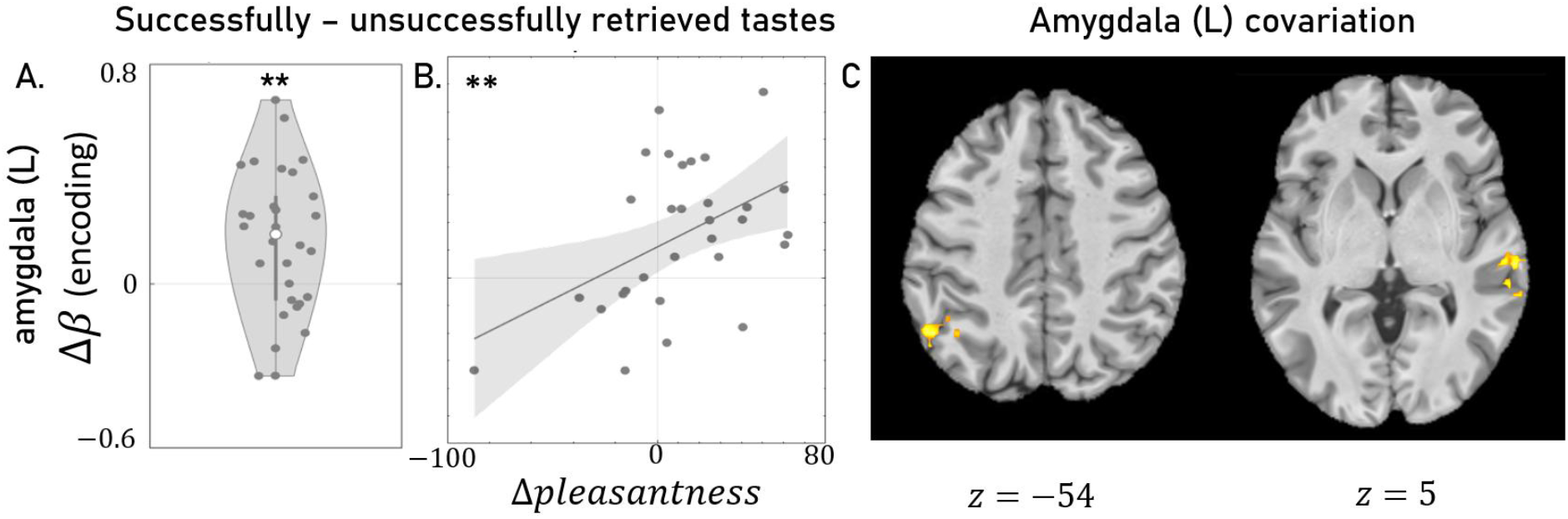
Amygdala activity during encoding predicts which cue-taste associations are retained. **(A) Encoding activity predicts memory retrieval.** An ROI search revealed encoding activity in the left amygdala that is higher during the encoding of successfully retrieved cue-taste pairs, compared with pairs retrieved less reliably. Because each pair was presented six times at encoding and probed repeatedly at retrieval, this contrast compares pairs by the reliability of later retrieval rather than by the fate of a single encoding event. **(B) Memory effect covaries with perceived pleasantness**. Across participants, the size of the amygdala effect covaried with a behavioral difference computed from the same partition: the difference in rated pleasantness between consistently and less reliably retrieved tastes. Each point is one participant; line shows the least-squares fit. **(C) Retrieval activity covaries with lateral parietal cortex.** Generalized psychophysiological interaction analysis with the left amygdala as seed. During taste retrieval trials, compared with visual control trials, amygdala activity covaried with left angular gyrus and a right cluster bordering the supramarginal gyrus and superior temporal sulcus. Amygdala ROI defined anatomically. Whole-brain analyses cluster-corrected at 2.3; no other region survived correction for the subsequent memory contrast. n = 30 (A, B), n=31 (C,D). Full cluster listings are given in Table S1.

Since pleasantness varies across tastes and the amygdala is sensitive to valence, we asked whether the two were related. For each participant we computed the difference in rated pleasantness between tastes later remembered consistently and those remembered less reliably, applying the same partition used for the neural analysis, and correlated this with the magnitude of the amygdala effect. The two covaried across participants (r(29) = .47, p<0.01; **Fig. 6B**): participants whose remembered tastes were the more pleasant ones showed larger amygdala differentiation at encoding.

We next asked whether the amygdala acts alone or as part of a network at retrieval, using a generalized psychophysiological interaction analysis with the amygdala as seed. Relative to visual control trials, the left amygdala showed increased covariation during taste retrieval with left angular gyrus and with a right lateral cluster spanning superior temporal sulcus and supramarginal gyrus (**Fig. 6C**) – the same lateral parietal regions in which encoding patterns generalized to retrieval. Against a resting baseline, coupling was also observed with left anterior PFC and gustatory cortex and with bilateral fusiform gyrus (Supplementary Fig. S4).

The amygdala’s contribution at encoding is therefore indicative of whether an association is retained. Its association with pleasantness is consistent with a role in registering the significance of an association rather than its sensory detail. At retrieval, however, it was coupled with the lateral parietal regions in which taste representation generalized from tasting to retrieval, suggesting that this contribution is not confined to the moment of learning.

## Discussion

What is recovered when a taste is retrieved? We asked which parts of the brain are active when a taste is remembered, and whether the representation at retrieval is the one the brain used while tasting. The gustatory cortex was activated when participants remembered a taste from a visual cue, even in the absence of gustatory input, and activity patterns in the gustatory cortex carried information about which taste was retrieved. But the specific patterns in the gustatory cortex established while a taste was in the mouth did not generalize to retrieval. Instead, patterns that generalized between experiencing a taste and later remembering it were found in lateral parietal cortex. Activity in the amygdala covaried with these same regions in the lateral parietal cortex during retrieval, while during encoding amygdala activation predicted which associations were most strongly retained in memory.

Recruitment of the gustatory cortex when a taste is remembered is itself a novel observation in humans. In this study, retrieval involved no gustatory stimulus of any kind: participants saw a bottle and reported which taste it was associated with in memory. Compared with a control condition matched for visual input, response demand, and the same images, the gustatory cortex responded more robustly when participants retrieved a taste. Moreover, activity patterns in gustatory cortex distinguished among specific tastes during retrieval, so its activity reflected not only engagement but information about which taste was being recalled.

These observations are consistent with a substantial body of work. Human imaging has shown gustatory cortex responses to food images, to taste imagery, and to cues that predict tastes^2,4–6^. Rodent electrophysiology has established that gustatory cortex is shaped by learning and participates directly in the formation and expression of cue-taste associations^7–10,14^. Our results extend these findings to memory of a specific taste, evoked by an arbitrary visual cue, learned within a single session.

Evidence of this kind has generally been read as indicating that gustatory cortex stores taste memories and that recalling a taste involves re-creating its sensory representation. Our findings show that the second part of this inference does not follow from the first. A region can be engaged at retrieval, and can carry stimulus-specific information there, while using a representation that is different from the one it used during perception. Distinguishing between reactivation and reinstatement requires asking whether a pattern learned in one phase generalizes to the other – a question that has not been asked in the gustatory domain.

The distinction between reactivation and reinstatement was not available to the taste literature, but it has been shown in vision and audition. Early work in those modalities showed that retrieving a memory reactivates cortical regions engaged during perception^16,34–36^. But as multivariate methods were applied to the question, it became clear that reactivation and reinstatement are not the same thing^21,23,25,33,37^. Applying that distinction to taste, we found that taste identity was decodable from the gustatory cortex during retrieval, but that the code was not the same as during tasting. The accuracy drop from encoding to cross-phase decoding was significantly larger in gustatory cortex than in lateral parietal cortex, and a search across the whole cortex found cross-phase generalization in frontal and parietal regions but none in gustatory cortex. The gustatory cortex therefore participates in remembering a taste without replaying the pattern it produced while tasting one.

The representation that did bridge the two phases was located outside the gustatory system. Patterns established while a taste was in the mouth transferred to recall in angular gyrus, medial frontal cortex, OFC and middle temporal cortex. Notably, the angular gyrus cluster identified by an unbiased whole-cortex search fell within a region independently specified on the basis of the visual memory literature, where retrieval-related representations are consistently found to be more abstract than their perceptual counterparts and are localized to parietal cortex^21,24–26^.

What survived the transition from experience to memory was not only the gist. The representation that transferred to lateral parietal cortex distinguished among individual tastes, not merely between categories; so, what the brain carried forward from tasting included which taste had been experienced, not only its category or valence. This sets the present results apart from an account in which memory preserves category and discards detail: the representation changed in format, but it did not simply coarsen. Nor was it only valence. Taste quality is tied to approach and avoidance through circuits that do not require learning^38–40^, which raises the possibility that recalling a taste might depend on reactivating those circuits rather than on recovering anything specific to the taste itself.

Memory for a taste was not all-or-none. Participants’ choices revealed that their memory often recovered the specific taste a cue predicted, but that even when they did not, they still remembered the taste category. Both levels of the memory were therefore available, and they could be separated. The brain data show the same separability. At retrieval, patterns in medial frontal cortex distinguished sweet from sour but did not discriminate individual tastes. These medial regions have been implicated in goal-directed behavior including value-guided decisions involving food^41–45^, and decisions of this kind depend more on whether a taste will be sweet or sour than on which sweet or sour taste it is. Category and identity are thus not one representation at two resolutions but distinguishable states, present in different places.

Category and identity also differed in how they crossed between phases. Identity transferred from tasting to recall in angular gyrus whereas category showed only weak transfer in the same region, but showed more robust transfer in other frontal and association cortical areas. Neither level transferred in gustatory cortex. Still, gustatory cortex patterns at retrieval carried both taste category and identity, consistent with accounts that treat the gustatory cortex as an integrative sensory region representing more than taste quality alone^31,32,46^.

Having two levels of specificity, in behavior and in the brain, is what makes questions about representation stability tractable. A transformed memory may have lost detail, kept it, or reorganized it. Distinguishing these requires a measure of specificity on both sides. In this case, what was coded in parietal cortex retained item level information, while what participants could report often did not.

Memory also varied across pairs and across people. Some associations were retained reliably and others were not, and this variability allows a different question: not what a taste memory is made of, but what determines whether one is formed at all. Activity in the amygdala during encoding predicted which associations survived. Pairs that were later retrieved consistently had elicited greater amygdala activity when the taste was experienced and no other region showed this relationship. The effect also tracked pleasantness: participants for whom the remembered tastes were the more pleasant ones showed the largest difference in amygdala activity at encoding. What the amygdala registered, on this account, was not which taste was present but how much it mattered.

This connects two literatures on amygdala function that have largely developed apart. In rodents, studies of cue–taste and cue–reward learning has established the amygdala as a substrate of associative learning, encoding the behavioral significance of sensory stimuli through its connections with gustatory cortex, thalamus and the ventral tegmental area^8,10–12,14,47–52^. Human imaging studies have implicated the amygdala in emotional and reinforcement learning ^53,54^, but human work on taste has largely treated it as a valence region, concerned with pleasantness and value^13,55–57^. Our results show that amygdala activity in humans was associated with the formation of arbitrary cue–taste associations, and predicted which of them was retained. At retrieval, amygdala was coupled with the lateral parietal regions in which the encoding was recoverable. Its contribution may therefore extend beyond the moment of learning though our data do not establish what the coupling carries.

Three limitations bear on how these findings should be read. We found no generalization in the gustatory cortex and while this was verified across converging analyses – an ROI analysis across a range of feature counts, and an unbiased search across cortex that identified other regions – it is still a null effect. A spatially restricted effect below the resolution of these analyses can not be excluded. Second, angular gyrus supports semantic as well as episodic retrieval, and participants may have encoded each association partly as a label; that a classifier trained on one sweet and one sour taste generalized to tastes it had never seen argues against a purely verbal account, as does the reactivation in sensory regions, including gustatory cortex. But these do not rule out semantic contributions. This is less a confound than a candidate mechanism: a representation that is semantic in part is one kind of transformed representation. Third, we study one modality. Behavioral studies suggest that retrieval of a taste may be different from that of sights or sounds. Whether taste transforms more than vision, as the gustatory hierarchy and oral and interoceptive components of taste might predict, requires testing the same participants across modalities, and remains open.

Most decisions about what to eat are made before anything is tasted. A cue is seen, a taste is anticipated and the anticipation guides the choice. What our results suggest is that the taste recovered at that moment is not a simple replay of the sensory experience. The gustatory system is engaged – the gustatory cortex responds when a taste is remembered and carries information about which taste it is – but the representation it uses is not the one it used while tasting. The representation that bridges experience and memory is found, instead, in association cortex, where it preserves not only the gist of the taste but, which specific taste it was. That a system with a shallow sensory hierarchy nonetheless produces a memory representation held outside of sensory cortex suggests that this transformation is not a byproduct of hierarchical processing but a property of how experience becomes memory.

In vision, a memory and its perception can at least be compared side by side. In taste they cannot, and the consequence is that what guides a decision about food is never the taste itself but a representation built in its absence.

## Acknowledgements

We thank all members of the Shohamy lab for helpful discussions. We thank Tanya Tabachnik and the Advanced Instrumentation platform for the construction of an MRI compatible taste delivery system, as well as the Human Imaging platform at the Zuckerman Mind Brain Behavior Institute for help setting up this new testing configuration.

## Author contributions

S.B., A.L., I.K., and D.S. conceived the study, and designed the experiments.

S.B. and A.L. designed and constructed the taste delivery system.

S.B. and S.X. acquired all imaging data.

S.B. and D.S. wrote the manuscript with input from all authors.

S.B. performed all data analysis, and prepared figures.

## Declaration of generative AI and AI-assisted technologies in the manuscript preparation process

During the preparation of this work, the authors used Claude (Sonnet 4.6, Opus 5.5) in order to review and modify the decoding analyses, and for late stage editing of language. After using this tool/service, the author(s) reviewed and edited the content as needed and take(s) full responsibility for the content of the published article.

## Materials and Methods

### Participants

Thirty-seven healthy participants (19 females, mean age = 23.4 ± 4.2) were recruited through fliers posted on campus and the surrounding area in New York City. All experimental procedures were approved by the Institutional Review Board (IRB) at Columbia University and all participants provided informed consent before taking part in the study. Specific analyses required different exclusion criteria; the number of participants included in each analysis is reported below and in figure captions.

### Experimental procedure

Participants performed a cue-association task while being scanned with fMRI, consisting of 2 blocks of taste encoding and 3 blocks of memory retrieval. An additional memory test assessed each cue-taste memory outside of the scanner.

#### Stimuli

Six tastes were used, with the intention of including tastes with similar as well as different taste categories: apple juice and watermelon juice (sweet), unsweetened lemon juice and cranberry juice (sour) and unsweetened grapefruit juice and black tea (bitter). Each taste was randomly associated with 2 different visual cues, shaped as bottles. The twelve bottles were taken from an image dataset^58^.

#### Taste encoding

This phase consisted of “taste” trials intermixed with “rinse” trials. On *taste trials*, participants were presented with pairs of tastes and bottle images and instructed to learn “which potion comes in which bottle”. On each trial, a bottle appeared on the screen for 3.5 s, followed by taste delivery starting 0.5s after image onset and continuing for 3 s (0.5 ml).

Following each “taste” trial, a fixation cross appeared and then a rinse trial commenced. On *rinse trials*, a rinse cue appeared on the screen followed by water delivery (0.5 s after cue onset for 3 s, ∼0.5 ml). The taste encoding phase included 72 trials in total, over 2 blocks, each lasting 8.08 m. Each bottle-taste pair was presented 6 times, such that each taste was delivered 12 times.

#### Taste retrieval

This phase consisted of taste retrieval trials and visual control trials. On taste retrieval trials, participants viewed a cue (a droplet icon) indicating the trial type (1 s) followed by a presentation of a bottle image (3 s), and then a question (r 2 s) pertaining to the taste associated with the bottle (“Was the potion in the bottle sweet/sour/ bitter/fruity/smooth/refreshing?”). Participants replied by pressing one of three buttons, indicating yes/no/don’t recall.

Visual control trials followed similar timing, sequence, and response buttons, but asked participants to respond based on memory for the bottle’s appearance. On these trials, the cue was a bottle icon, followed by a question about the visual properties of the bottle (“Is the bottle pretty/blue/transparent?”).

Participants performed 108 trials overall, (90 taste retrieval trials, and 18 visual control trials, randomly intermixed), divided into 3 blocks, each lasting 8.37 m. Both taste encoding and retrieval used inter-trial interval jitters drawn from an exponential distribution.

#### Post scan taste assessment

At the end of the session, participants rated how sweet, sour, bitter and pleasant they found each taste, using a continuous sliding scale. They were then presented with all bottles used in the task and asked to choose the bottle that was associated with each taste. The first two participants did not perform the post scan assessment, yielding n=35 for all the post-scan related analyses.

### Setup

Tastants were delivered via a custom-built MRI-compatible system. Tastants were kept at room temperature in 250ml beakers connected to 12V Solenoid-Operated Pinch Valves. Fluid delivery was controlled by an arduino board that allowed the release of the tastants into polyurethane tubing that ran to a 3D printed mouthpiece^59^. The arduino board was connected to a stimulus delivery computer, which controlled the precise timing of tastants dispensed to the subject during the scan. Visual stimuli for behavioral and fMRI tasks were projected onto a screen located inside the scanner bore and viewed through a mirror system mounted on the head-coil. Both visual stimulus presentation and tastant delivery were controlled and synchronized via a custom-built program developed in the PsychoPy3 environment^60^.

### Image acquisition

fMRI data were collected at the MRI Platform at the Zuckerman Mind Brain Behavior Institute at Columbia University, using a 3T Siemens Priesma scanner and a 32-channel head coil. BOLD acquisition was done with epin-echo echoplanar imaging (SE-EPI) sequence (TE = 30 ms, TR = 1560 ms, flip angle = 65°, voxel size = 2 × 2 × 2 mm3, 72 axial slices). A multiband factor of 3 was used to acquire data from multiple slices simultaneously. A GRAPPA factor of 2 was used for in-plane slice acceleration. Each slice was oriented in the axial plane, with an anterior-to-posterior phase encoding direction. Before the task, 1 min EPI scans were acquired with opposite phase encoding direction (anterior-to-posterior and posterior-to-anterior), which were used for correction of spatial distortion artifacts during preprocessing. An additional high resolution MPRAGE sequence was used to provide an anatomical reference for the fMRI analysis (TE = 2.98 ms, TR = 2300 ms, flip angle = 9°, voxel size = 1 × 1 × 1 mm).

### Preprocessing

#### Preprocessing of B_0_ inhomogeneity mappings

A total of 1 fieldmaps were found available within the input BIDS structure for this particular subject. A *B_0_*-nonuniformity map (or *fieldmap*) was estimated based on two (or more) echo-planar imaging (EPI) references with topup^64^; FSL.

#### Anatomical data preprocessing

A total of 1 T1-weighted (T1w) images were found within the input BIDS dataset. The T1w image was corrected for intensity non-uniformity (INU) with N4BiasFieldCorrection^65^, distributed with ANTs 2.5.3^66^ (RRID:SCR_004757), and used as T1w-reference throughout the workflow. The T1w-reference was then skull-stripped with a *Nipype* implementation of the antsBrainExtraction.sh workflow (from ANTs), using OASIS30ANTs as target template. Brain tissue segmentation of cerebrospinal fluid (CSF), white-matter (WM) and gray-matter (GM) was performed on the brain-extracted T1w using fast (FSL (version unknown), RRID:SCR_002823^67^). Brain surfaces were reconstructed using recon-all (FreeSurfer 7.3.2, RRID:SCR_001847^68^), and the brain mask estimated previously was refined with a custom variation of the method to reconcile ANTs-derived and FreeSurfer-derived segmentations of the cortical gray-matter of Mindboggle (RRID:SCR_002438^69^). Volume-based spatial normalization to one standard space (MNI152NLin2009cAsym) was performed through nonlinear registration with antsRegistration (ANTs 2.5.3), using brain-extracted versions of both T1w reference and the T1w template. The following template was were selected for spatial normalization and accessed with *TemplateFlow* (24.2.0^70^): *ICBM 152 Nonlinear Asymmetrical template version 2009c*^71^ [RRID:SCR_008796; TemplateFlow ID: MNI152NLin2009cAsym].

#### Functional data preprocessing

For each of the 5 BOLD runs found per subject (across all tasks and sessions), the following preprocessing was performed. First, a reference volume was generated, using a custom methodology of *fMRIPrep*, for use in head motion correction. Head-motion parameters with respect to the BOLD reference (transformation matrices, and six corresponding rotation and translation parameters) are estimated before any spatiotemporal filtering using mcflirt (FSL^72^). The estimated *fieldmap* was then aligned with rigid-registration to the target EPI (echo-planar imaging) reference run. The field coefficients were mapped on to the reference EPI using the transform. The BOLD reference was then co-registered to the T1w reference using bbregister (FreeSurfer) which implements boundary-based registration^73^. Co-registration was configured with six degrees of freedom. Several confounding time-series were calculated based on the *preprocessed BOLD*: framewise displacement (FD), DVARS and three region-wise global signals. FD was computed using two formulations following Power (absolute sum of relative motions^74^) and Jenkinson (relative root mean square displacement between affines^72^). FD and DVARS are calculated for each functional run, both using their implementations in *Nipype* (following the definitions by Power et al. 2014^74^). The three global signals are extracted within the CSF, the WM, and the whole-brain masks. Additionally, a set of physiological regressors were extracted to allow for component-based noise correction (*CompCor*^75^). Principal components are estimated after high-pass filtering the *preprocessed BOLD* time-series (using a discrete cosine filter with 128s cut-off) for the two *CompCor* variants: temporal (tCompCor) and anatomical (aCompCor). tCompCor components are then calculated from the top 2% variable voxels within the brain mask. For aCompCor, three probabilistic masks (CSF, WM and combined CSF+WM) are generated in anatomical space. The implementation differs from that of Behzadi et al. in that instead of eroding the masks by 2 pixels on BOLD space, a mask of pixels that likely contain a volume fraction of GM is subtracted from the aCompCor masks. This mask is obtained by dilating a GM mask extracted from the FreeSurfer’s *aseg* segmentation, and it ensures components are not extracted from voxels containing a minimal fraction of GM. Finally, these masks are resampled into BOLD space and binarized by thresholding at 0.99 (as in the original implementation). Components are also calculated separately within the WM and CSF masks. For each CompCor decomposition, the *k* components with the largest singular values are retained, such that the retained components’ time series are sufficient to explain 50 percent of variance across the nuisance mask (CSF, WM, combined, or temporal). The remaining components are dropped from consideration. The head-motion estimates calculated in the correction step were also placed within the corresponding confounds file. The confound time series derived from head motion estimates and global signals were expanded with the inclusion of temporal derivatives and quadratic terms for each^76^. Frames that exceeded a threshold of 0.5 mm FD or 1.5 standardized DVARS were annotated as motion outliers. Additional nuisance timeseries are calculated by means of principal components analysis of the signal found within a thin band (*crown*) of voxels around the edge of the brain, as proposed by Patriat et al., (2017)^77^. All resamplings can be performed with *a single interpolation step* by composing all the pertinent transformations (i.e. head-motion transform matrices, susceptibility distortion correction when available, and co-registrations to anatomical and output spaces). Gridded (volumetric) resamplings were performed using nitransforms, configured with cubic B-spline interpolation.

Many internal operations of *fMRIPrep* use *Nilearn* 0.10.478 (RRID:SCR_001362), mostly within the functional processing workflow. For more details of the pipeline, see the section corresponding to workflows in *fMRIPrep*’s documentation (https://fmriprep.org/en/latest/workflows.html).

### Analysis

#### Univariate contrasts

Univariate analysis was performed using FSL^79^ in the framework of a general linear model (GLM). The model included 13 confounds as nuisance regressors (3 global signals variables and 10 motion variables) as well as task event onsets, convolved with the canonical hemodynamic response function and its time derivative. For taste encoding, the model included separate regressors for the onsets of the 6 different tastes, as well as a regressor for the rinse period. For the taste retrieval the model included 7 regressors for image onsets (6 for the taste-associate trials and 1 for the visual condition), as well as 2 regressors for the question period (1 per condition). At the group level, for both tasks the significance threshold was set to p < 0.05, family-wise error (FWE) corrected for multiple comparisons at the cluster level, with an underlying threshold of z > 2.3 at the peak level.

To examine taste-encoding related activity we contrasted the juice delivery period to the rinse period. For taste-retrieval we contrasted taste retrieval trials with visual-control trials. To identify regions involved in both taste encoding and taste retrieval, we conducted a conjunction analysis on the second-level z-statistic maps from each contrast. The conjunction map was computed as the voxel-wise minimum of the two z-statistic maps, following the minimum statistic approach^80^. Cluster-correction was performed using FSL with a cluster-forming threshold of z > 2.3 and a corrected cluster significance threshold of p < 0.05.

One participant was missing encoding onsets, and another participant chose to stop the scan before taste retrieval, resulting in thirty six participants included in the encoding and retrieval analyses.

#### Multivariate analyses: Taste decoding

We estimated trial-wise GLM beta estimates using GLMsingle^81^. GLMs for taste learning used 12 task-related regressors reflecting the onsets of the 12 different bottles. For retrieval, in addition to the 12 regressors reflecting each bottle’s appearance during taste-retrieval trials, we also included regressors for visual control trials, for cue appearance, and for button presses, resulting in 15 task-relevant regressors. Due to a design issue with the jitter of the intertrial interval, beta estimates could not be computed for the encoding of 3 participants, and the retrieval of 1 additional participant.

For taste identity decoding, we trained a classifier to discriminate each of the 6 tastes from the remaining 5 in a one-vs-rest framework. Each taste was associated with 2 bottles, and we used a leave-one-bottle-out cross-validation scheme, yielding 12 folds total (2 folds × 6 tastes). Critically, on each fold, the held-out bottle of the target taste was tested against the held-out bottle of every other taste, ensuring balanced train and test set composition and avoiding any bottle-position confound. For taste category decoding we trained a classifier to discriminate between sweet and sour tastes (watermelon and apple, compared with lemon and cranberry). Using a leave-one-taste-out cross-validation scheme, provided 4 folds: each fold trains on one sweet taste + one sour taste (both bottles of each) and tests on the complementary sweet + sour taste (both bottles of each), cycling through all 4 sweet–sour pairings. In both decoding approaches we used a linear support vector classifier (SVC, C = 0.1). To account for class imbalance inherent to the one-vs-rest design, the classifier was trained with balanced class weights, and performance was evaluated using balanced accuracy. The final decoding accuracy per subject was taken as the mean balanced accuracy across all valid folds, with chance level at 50%. Features were z-scored prior to classification using a standard scaler fit on the training set. The analysis was implemented in Python (v3.10.14) using the scikit-learn package (v1.4.2^82^).

We only performed this analysis on subjects who had both encoding and retrieval single trial beta estimates (n=32). Additionally, we excluded retrieval trials on which subjects reported not recalling the correct response. Since we required every decoding group to have at least 4 trials available for training, and included only subjects for whom every group met this criterion (4 groups for taste category, 12 for flavor identity), 4 subjects were excluded from the within-retrieval taste identity decoding (n=28 included in the analysis), and one subject excluded from within retrieval decoding of taste category (n=31 included in the analysis). We retained these subjects in the cross-phase (encoding-to-retrieval) decoding, since their encoding data still provided sufficient trials for training.

The classifiers were applied to the whole brain, using a surface-based Searchlight approach. First, we projected the single-trial beta estimates to the fsaverage6 template. For each vertex on the cortical surface, a local neighborhood of vertices was defined using a precomputed searchlight list, and the MVPA pipeline described above was applied to the beta patterns within that searchlight. Searchlights with fewer than 50 vertices after quality screening were excluded. Vertex-wise beta estimates were cleaned prior to decoding by removing vertices whose mean or standard deviation across trials was a statistical outlier (MAD-based, threshold = 3). The resulting accuracy values were projected back onto the surface using a Gaussian-weighted accumulation scheme, in which each vertex’s contribution was weighted by its distance from the searchlight center (σ = 0.5 × mean distance). This yielded a smoothed, vertex-wise accuracy map for each participant. Group-level inference was performed using a one-sample t-test against the null hypothesis (50%). To correct for multiple comparisons, we applied a non-parametric permutation-based cluster-level FWE correction (5000 permutations). For the encoding stage, where the effects were strong, we use a voxel threshold of p<0.001, and for retrieval p<0.05. For both stages we used the same cluster threshold of p<0.05. For visualization purposes, z-statistics within surviving clusters were smoothed across neighboring vertices (5 iterations of surface-based neighbor averaging).

The classifiers were also applied to 2 ROIs: the gustatory cortex and the angular gyrus. These ROIs were chosen based on previous literature implicating them in taste perception and memory reinstatement, respectively. The ROIs were defined in cortical surface space using the Destrieux atlas (aparc.a2009s). The masks were extracted in fsaverage5 resolution (10,242 vertices/hemisphere), and resampled to fsaverage6 using nearest-neighbor interpolation. The gustatory cortex ROI was formed using the union of the short insular gyri and the anterior and superior circular insular sulci. To characterize decoding strength in each surface ROI without committing to a single, arbitrarily-chosen feature count, we ran the SVM decoder across a range of feature counts for each ROI, hemisphere, subject, and condition: k was set to six fixed percentages (2%, 5%, 10%, 25%, 50%, 80%) of that ROI’s own cleaned vertex count (features selected by univariate F-test, SelectKBest), yielding a permutation-derived z-score at each of the six feature counts. We summarized each subject’s full z-score-vs-log10(k) curve as a single area-under-the-curve (AUC) value, computed by trapezoidal integration and normalized by the log10(k) range swept. This yields a quantity in the same units as the underlying z-scores: the average decoding strength a subject would show if the effect held constant across the tested feature-count range. For calculating statistical significance, we averaged participants’ left and right hemisphere, as we did not have a-priori reason to expect laterality. Group-level significance for each ROI × task × phase condition was assessed with a one-sample t-test on subjects’ AUC values against zero. We further used a linear model to directly test for an interaction between the ROIs and the learning phase (encoding compared with cross-phase decoding).

#### Subsequent memory effects

We defined memory performance using memory retrieval trials, comparing pairs that were later remembered perfectly (100% accuracy) vs. those for which memory was weaker (<100%). This measure of accuracy was calculated over a subset of trials: on some trials, accuracy could not be defined because the nature of the memory question was ambiguous with regards to the perception (e.g. was the taste fruity?), whereas questions about whether the test was sweet, bitter, or sour were ones for which we could verify the participant’s subjective perception, giving us confidence in rating their memory performance. Each taste had 7 such trials on average per person (ranging from 2 to 12 trials).

Using this sorting measure, for each participant we looked at the difference in activity between tastes that were perfectly remembered and those that included mistakes. The activity difference was calculated using the trial-wise beta estimates during taste encoding. We used a t-test to assess whether encoding activity was greater for successfully retrieved tastes than those that were not successfully retrieved. We calculated the difference in the amygdala and the hippocampus, defined using the Tian subcortical atlas^83^, two a-priori ROIs. Additionally, we performed a whole brain analysis using 300 cortical parcels^84^ and 54 subcortical regions^83^, and calculated adjusted p-values using the Benjamini-Hochberg procedure to control the false discovery rate^85^. This analysis included participants with trial-level beta estimates at encoding, and excluded a participant that perfectly remembered all tastes, and 2 that did not remember any of the tastes (n=30).

#### Task-based functional connectivity

We found the amygdala activity at encoding is greater for tastes that were better remembered. As a post-hoc analysis, designed to evaluate the amygdala’s role during retrieval, we used generalized psychophysiological interactions (gPPI), implemented with FSL. gPPI estimates functional connectivity by constructing an interaction term between the time-course in a seed region of interest and a task regressor. Voxels whose activity are well fit by this interaction term are assumed to be “functionally coupled” with the seed region in a way that depends on the behavioral task^86,87^. To calculate the amygdala’s psychophysiological interactions during taste retrieval, the left amygdala ROI was defined using FreeSurfer parcellation, and its time series was extracted using fslmeants, and included as a regressor in a GLM. The model also included nuisance regressors (identical to the original univariate analysis), and task-regressors, namely the onset of images for taste retrieval trials, the onsets of images for visual control trials, and the onsets of images in missed trials. We then examined the interaction between the “taste retrieval trial” coefficient and the areas covarying with the amygdala. Like other connectivity measures, gPPI depends on covariance between regional time series, which head motion is known to inflate^88^, including in task-based designs^89^. Therefore, we excluded 5 participants exhibiting the highest amount of motion (> 10% of volumes were labeled as motion outliers), (n=31).

