## Supplemental Information for "Tasting from memory: reconstructing taste from visual cues in the human brain"

**Supplementary Information**

**Supplementary Figure S1. Taste identity and category are widely decodable during encoding. (A) Taste identity** was decodable across the frontal lobe and in the temporal, parietal, and occipital lobes. **(B) Taste category** was decodable across most of the cortex.

Whole brain maps computed with a surface-based searchlight. Results are projected on the fsaverage6 surface. Color indicates z-statistics from permutation testing against chance (1/6 for identity, 1/2 for category). Maps were cluster-corrected using permutation-based cluster-level familywise error correction with a voxel threshold of $p<0.05$ and smoothed for visualization; statistics were computed on unsmoothed data. The corresponding encoding maps at a threshold of $p<0.001$ are presented in the main Figs 3 and 5.

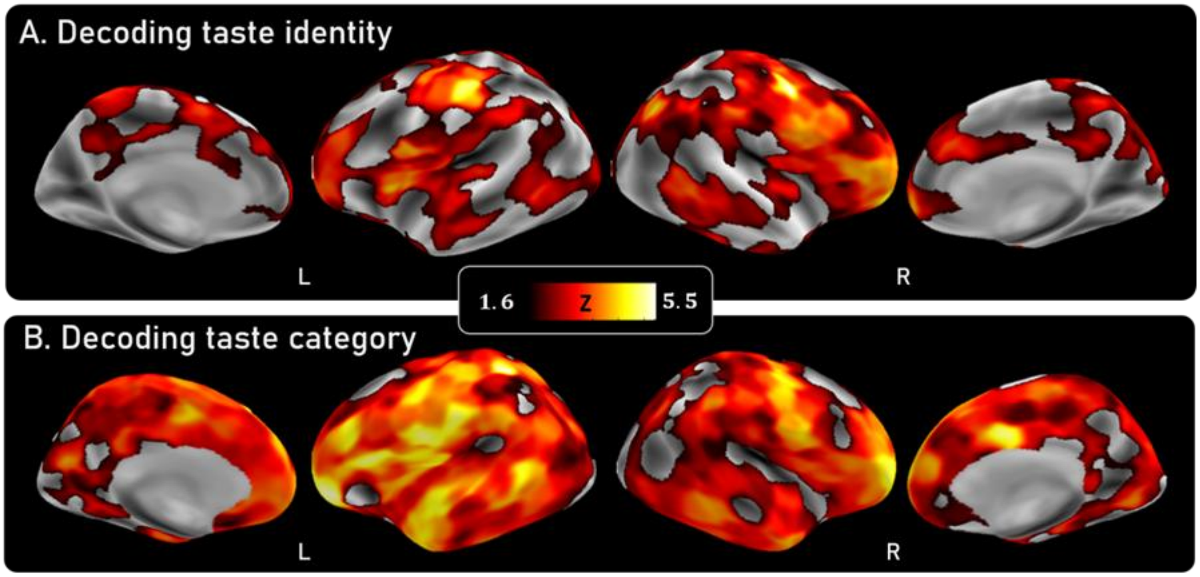

**Supplementary Figure S2. ROI decoding dependency on feature selection.** Decoding accuracy in gustatory cortex and lateral parietal cortex across 6 feature selection thresholds, defined as percent of the ROI vertices. **A. Gustatory cortex,** for taste identity (top row) and taste category (bottom row) at each task phase. During encoding (left), decoding accuracy was high at any feature selection threshold. Decoding accuracy varied as a function of feature selection threshold at retrieval (middle), and was around chance for cross-phase decoding (right). **B. Lateral parietal cortex,** for taste identity (top row) and taste category (bottom row). Decoding accuracy was high at all feature selection thresholds during encoding. During retrieval, decoding varied as a function of feature selection thresholds. Cross-phase decoding of taste identity similarly varies, but this variation only reaches significance for taste identity decoding.
Filled symbols indicate $p < .05$; error bars show SEM. The trapezoidal integration of the decoding accuracy across the feature space is presented in the main text (**Fig. 4**), and in supplementary Fig. S3.

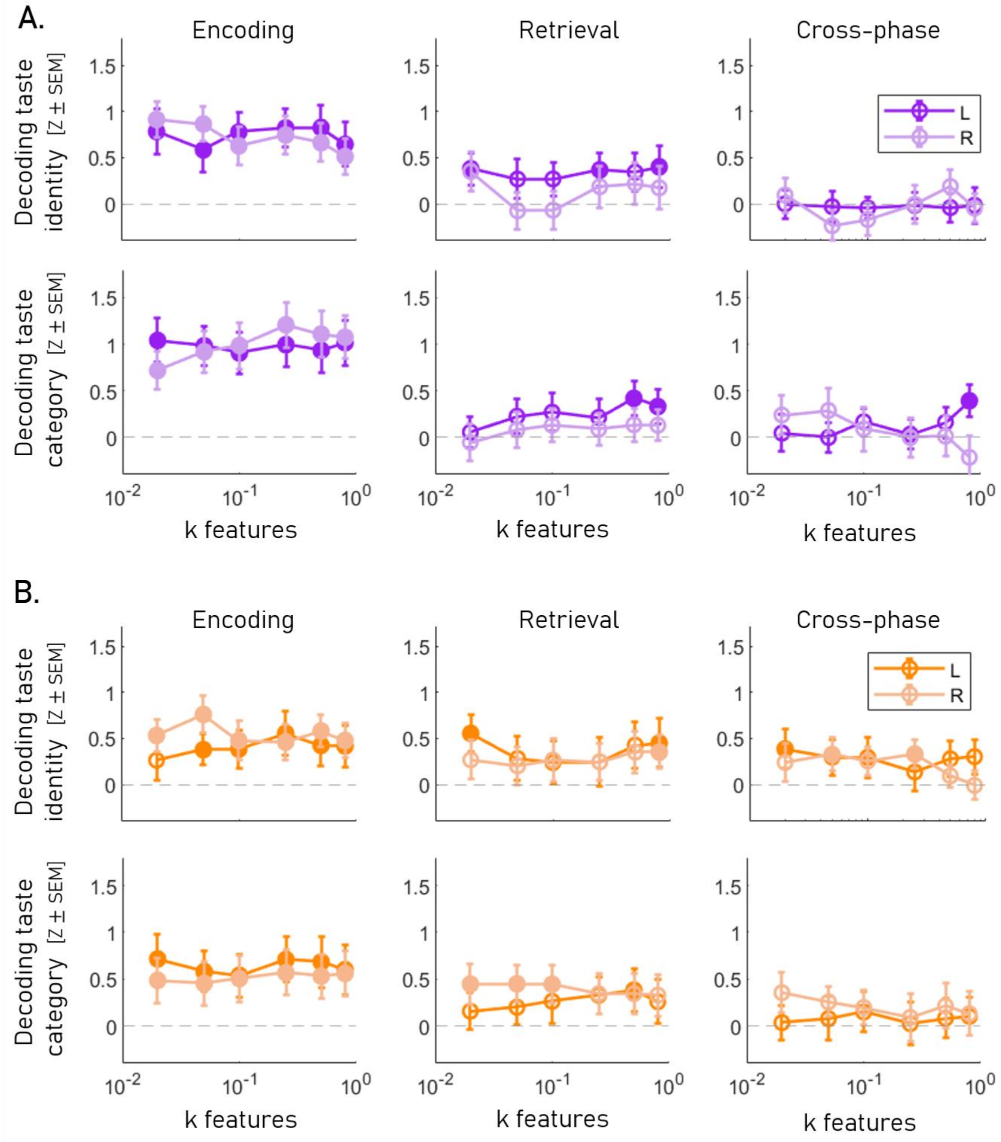

**Supplementary Figure S3. ROI taste category decoding.** Decoding taste category in gustatory cortex (anterior dorsal insula, purple) and lateral parietal cortex (angular gyrus, orange), defined anatomically and specified in advance, shown separately for each hemisphere. The accuracy is plotted for encoding, retrieval and cross-phase decoding. Both ROIs represented taste category during encoding. Gustatory cortex showed no evidence of cross-phase generalization. Lateral parietal cortex represented taste category during retrieval but did not display generalization across learning phases. Filled symbols indicate p < .05; error bars show SEM. The ROIs are displayed below, on the fsaverage6 inflated brain.

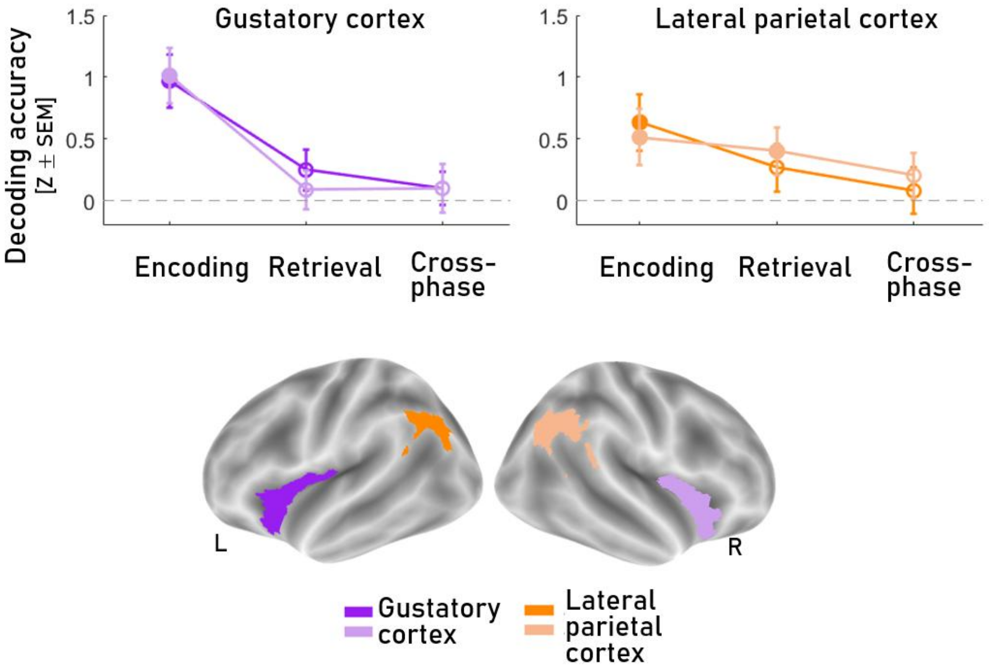

**Supplementary Figure S4. Amygdala connectivity during taste retrieval**. Generalized psychophysiological interaction analysis with the left amygdala as seed. Relative to baseline, taste retrieval trials were associated with greater functional coupling between left amygdala and left gustatory cortex, fusiform gyrus, and anterior PFC. No taste was delivered during this phase. Whole-brain analysis, cluster-corrected at z>2.3; n=31; Full cluster listings are given in Table S1.

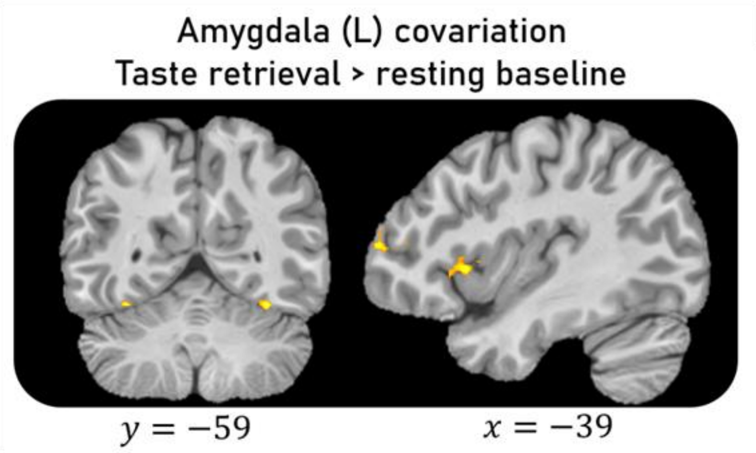

.

**Table S1: Cluster listings for univariate and connectivity analyses.** Clusters from each whole-brain contrast, listed by section: encoding phase (taste>rinse), retrieval (taste retrieval>visual control), and conjunction contrasts, shown in main Fig. 2, and generalized psychophysiological interaction analyses with the left amygdala as seed, contrasted against visual control trials (main Fig 6) and against baseline (Supp Fig. 3). For each cluster: extent in voxels; the anatomical label at the peak coordinate, with *probabilistic Harvard-Oxford atlas confidence in parentheses; hemisphere; peak z; and peak MNI coordinates. The final column lists* additional *regions accounting for at least 5% of the cluster's volume, with* the *percentage of* cluster *volume in each. Peaks use a cluster-forming threshold of z>2.3.*

| **Encoding** | | | | | | | |
|---|---|---|---|---|---|---|---|
| **peak** | | | | | | | |
| **Region** | **Hemi** | **Z** | **MNI x** | **MNI y** | **MNI z** | **Extent (voxels)** | **Other labels >=5% (atlas)** |
| Inferior Temporal Gyrus, posterior division (16%) | Right | 7.92 | 44 | -22 | -36 | 27311 | PosteriorTemporal Fusiform Cortex (6%) |
| Juxtapositional Lobule Cortex (formerly Supplementary Motor Cortex) (78%) | Left | 5.26 | -4 | -4 | 64 | 2135 | |
| Postcentral Gyrus (35%) | Left | 3.85 | -58 | -24 | 24 | 304 | Anterior Supramarginal (33%); Central Opercular Cortex (8%); Parietal Operculum Cortex (5%) |
| **Retrieval** | | | | | | | |
| **peak** | | | | | | | |
| **Region** | **Hemi** | **Z** | **MNI x** | **MNI y** | **MNI z** | **Extent (voxels)** | **Other labels >=5% (atlas)** |
| Juxtapositional Lobule Cortex (formerly Supplementary Motor Cortex) (48%) | Left | 6.83 | -4 | 8 | 58 | 31735 | Superior Frontal Gyrus (29%); Paracingulate Gyrus (5%) |
| Insular Cortex (48%) | Right | 5.18 | 40 | 14 | 2 | 1412 | Frontal Operculum Cortex (30%); Central Opercular Cortex (5%) |
| Supramarginal Gyrus, posterior division (15%) | Right | 4.24 | 58 | -38 | 10 | 727 | TemporooccipitaMiddle Temporal Gyrus (12%); posterior Superior Temporal Gyrus (7%); PosteriorMiddle Temporal Gyrus (5%) |
| Cingulate Gyrus, posterior division (32%) | Left | 4.33 | -6 | -38 | 26 | 299 | |
| Frontal Pole (81%) | Left | 4.13 | -30 | 44 | 32 | 295 | Middle Frontal Gyrus (5%) |
| Left VIIIa (76%, Cerebellum (MNIflirt)) | Left | 4.25 | -24 | -62 | -50 | 270 | Left VIIb (Cerebellum (MNIflirt)) (14%) |
| Frontal Pole (89%) | Right | 3.74 | 28 | 46 | 36 | 220 | |
| Lateral Occipital Cortex, inferior division (57%) | Right | 3.68 | 46 | -72 | 6 | 172 | |

| Conjunction | | | | | | | |
|---|---|---|---|---|---|---|---|
| peak | | | | | | | |
| Region | Hemi | Z | MNI x | MNI y | MNI z | Extent (voxels) | Other labels >=5% (atlas) |
| Juxtapositional Lobule Cortex (formerly Supplementary Motor Cortex) (78%) | Left | 5.26 | -4 | -4 | 64 | 1204 | |
| Insular Cortex (43%) | Right | 4.89 | 30 | 16 | 8 | 742 | |
| Brain-Stem (51%) | Left | 4.53 | -4 | -28 | -6 | 721 | |
| Occipital Fusiform Gyrus (56%) | Left | 5.04 | -30 | -80 | -14 | 652 | Inferior Lateral Occipital Cortex (15%) |
| Insular Cortex (72%) | Left | 4.55 | -30 | 22 | 6 | 476 | Frontal Operculum Cortex (7%) |
| Occipital Fusiform Gyrus (62%) | Right | 4.63 | 24 | -78 | -12 | 474 | Lingual Gyrus (7%) |
| **gPPI: Left amygdala X taste memory - visual control** | | | | | | | |
| peak | | | | | | | |
| Region | Hemi | Z | MNI x | MNI y | MNI z | Extent (voxels) | Other labels >=5% (atlas) |
| Angular Gyrus (50%) | Left | 4.15 | -52 | -58 | 46 | 272 | Superior Lateral Occipital Cortex (29%); posterior Supramarginal Gyrus (11%) |
| Supramarginal Gyrus, posterior division (22%) | Right | 4.15 | 64 | -36 | 6 | 216 | Posterioe Superior Temporal Gyrus (17%); posterior Middle Temporal Gyrus (9%); termporoocipital Middle Temporal Gyrus (5%) |
| **gPPI: Left amygdala X taste memory** | | | | | | | |
| peak | | | | | | | |
| Region | Hemi | Z | MNI x | MNI y | MNI z | Extent (voxels) | Other labels >=5% (atlas) |
| Frontal Pole (42%) | Left | 3.5 | -36 | 60 | 12 | 107 | |
| Frontal Operculum Cortex (61%) | Left | 4.24 | -44 | 16 | 0 | 126 | Inferior Frontal Gyrus, pars opercularis (7%); Inferior Frontal Gyrus, pars triangularis (5%) |
| Temporal Occipital Fusiform Cortex (40%) | Right | 3.98 | 34 | -62 | -18 | 121 | Occipital Fusiform Gyrus (33%); Right VI (11%) (Cerebellum (MNIflirt)) |
| Temporal Occipital Fusiform Cortex (64%) | Left | 3.9 | -34 | -60 | -18 | 107 | Occipital Fusiform Gyrus (18%) |
| Occipital Pole (60%) | Left | 3.86 | -30 | -96 | -8 | 141 | Inferior Lateral Occipital Cortex (10%) |

**Table S2: Cluster listings for taste identity decoding.** For each cluster from the encoding, and retrieval searchlight analyses (main Fig. 3): cluster ID, mean decoding accuracy (chance=1/6), hemisphere, and the anatomical regions it spans. Regions are given as Destrieux atlas parcels (aparc.a2009s), grouped under broader anatomical headings for readability. For each parcel, the percentage of the cluster those vertices represent, and the percentage of the parcel falling within the cluster. Parcels are listed if they account for at least 5% of the cluster or if at least 50% of the parcel falls within the cluster.

| Encoding | | | | | | | |
|---|---|---|---|---|---|---|---|
| main areas (R) | hemi | subregions | | | | accuracy (mean) | cluster id |
| | | dk region | vertices | % of cluster | % of region | | |
| Premotor / precentral | Right | G_precentral | 528 | 13.6 | 56.9 | 0.52 | 1 |
| | Right | S_precentral-inf-part | 372 | 9.6 | 78.5 | 0.52 | 1 |
| | Right | G_front_middle | 312 | 8.0 | 30.7 | 0.52 | 1 |
| Ventrolateral prefrontal | Right | S_front_inf | 301 | 7.7 | 51.3 | 0.52 | 1 |
| | Right | S_front_middle | 238 | 6.1 | 46.8 | 0.52 | 1 |
| | Right | G_front_inf-Triangul | 221 | 5.7 | 80.4 | 0.52 | 1 |
| | Right | G_and_S_frontomargin | 191 | 4.9 | 87.6 | 0.52 | 1 |
| Orbitofrontal / frontopolar | Right | G_and_S_transv_frontopol | 199 | 5.1 | 90.9 | 0.52 | 1 |
| | Right | G_orbital | 206 | 5.3 | 30.7 | 0.52 | 1 |
| | Right | S_orbital_lateral | 97 | 2.5 | 100.0 | 0.52 | 1 |
| | Right | G_front_inf-Orbital | 68 | 1.7 | 77.3 | 0.52 | 1 |
| Peri-central | Left | S_central | 421 | 33.8 | 35.1 | 0.516 | 2 |
| | Left | G_postcentral | 391 | 31.4 | 43.6 | 0.516 | 2 |
| | Left | G_precentral | 191 | 15.3 | 19.3 | 0.516 | 2 |
| | Left | S_postcentral | 143 | 11.5 | 13.1 | 0.516 | 2 |
| | Left | S_precentral-sup-part | 83 | 6.7 | 18.2 | 0.516 | 2 |
| Operculo-insular | Left | S_circular_insula_sup | 446 | 73.2 | 59.1 | 0.518 | 3 |
| | Left | G_Ins_lg_and_S_cent_ins | 61 | 10.0 | 29.2 | 0.518 | 3 |
| | Left | G_insular_short | 47 | 7.7 | 20.8 | 0.518 | 3 |
| Orbitofrontal / frontopolar | Left | G_front_middle | 228 | 26.6 | 21.8 | 0.514 | 4 |
| | Left | G_and_S_frontomargin | 147 | 17.2 | 62.8 | 0.514 | 4 |
| | Left | S_front_inf | 133 | 15.5 | 23.5 | 0.514 | 4 |
| | Left | G_orbital | 112 | 13.1 | 18.2 | 0.514 | 4 |
| | Left | S_orbital_lateral | 77 | 9.0 | 98.7 | 0.514 | 4 |
| | Left | G_and_S_transv_frontopol | 76 | 8.9 | 52.4 | 0.514 | 4 |

| Retrieval | | | | | | | |
|---|---|---|---|---|---|---|---|
| main areas (R) | hemi | subregions | | | | accuracy (mean) | cluster id |
| | | dk region | vertices | % of cluster | % of region | | |
| Parieto-occipital | Right | G_pariet_inf-Angular | 285 | 78.1 | 29.8 | 0.51 | 1 |
| | Right | S_intrapariet_and_P_trans | 70 | 19.2 | 6.1 | 0.51 | 1 |
| Temporo-occpital | Right | G_and_S_occipital_inf | 212 | 50.8 | 79.7 | 0.51 | 2 |
| | Right | G_temporal_inf | 72 | 17.3 | 13.0 | 0.51 | 2 |
| | Right | S_oc-temp_lat | 66 | 15.8 | 26.1 | 0.51 | 2 |
| | Right | Pole_occipital | 40 | 9.6 | 5.9 | 0.51 | 2 |
| Temporal | Left | S_temporal_sup | 600 | 58.9 | 35.6 | 0.51 | 3 |
| | Left | G_temporal_middle | 253 | 24.8 | 36.4 | 0.51 | 3 |
| | Left | G_temporal_inf | 63 | 6.2 | 9.7 | 0.51 | 3 |
| | Left | S_oc-temp_lat | 56 | 5.5 | 24.2 | 0.51 | 3 |
| Posterior cingulate | Right | S_subparietal | 163 | 52.9 | 32.1 | 0.51 | 4 |
| | Right | G_cingul-Post-dorsal | 93 | 30.2 | 44.7 | 0.51 | 4 |
| | Right | S_pericallosal | 45 | 14.6 | 8.0 | 0.51 | 4 |
| Insula | Right | S_circular_insula_inf | 162 | 44.9 | 33.0 | 0.51 | 5 |
| | Right | G_Ins_lg_and_S_cent_ins | 94 | 26.0 | 43.3 | 0.51 | 5 |
| | Right | G_insular_short | 46 | 12.7 | 22.1 | 0.51 | 5 |
| | Right | G_temp_sup-Plan_polar | 39 | 10.8 | 17.8 | 0.51 | 5 |
| Anterior cingulate / superior frontal | Right | G_front_sup | 285 | 64.2 | 14.5 | 0.51 | 6 |
| | Right | G_and_S_cingul-Ant | 99 | 22.3 | 13.1 | 0.51 | 6 |
| | Right | S_front_sup | 57 | 12.8 | 7.5 | 0.51 | 6 |
| Superior frontal | Left | S_front_sup | 302 | 27.9 | 38.4 | 0.51 | 7 |
| | Left | S_front_middle | 284 | 26.2 | 76.1 | 0.51 | 7 |
| | Left | G_and_S_frontomargin | 165 | 15.2 | 70.5 | 0.51 | 7 |
| | Left | G_front_sup | 133 | 12.3 | 6.6 | 0.51 | 7 |
| | Left | G_front_middle | 114 | 10.5 | 10.9 | 0.51 | 7 |
| Precentral | Left | G_precentral | 269 | 61.6 | 27.2 | 0.51 | 8 |
| | Left | S_precentral-inf-part | 156 | 35.7 | 32.0 | 0.51 | 8 |

**Table S3: Cluster listings for taste identity cross-phase decoding.** For each cluster from the cross-phase taste identity searchlight analysis (main Fig. 4); cluster ID, mean decoding accuracy (chance=1/6), hemisphere, and the anatomical regions it spans. Regions are given as Destrieux atlas parcels (aparc.a2009s), grouped under broader anatomical headings for readability. For each parcel, the percentage of the cluster those vertices represent, and the percentage of the parcel falling within the cluster. Parcels are listed if they account for at least 5% of the cluster or if at least 50% of the parcel falls within the cluster.

| Taste identity | | | | | | | |
|---|---|---|---|---|---|---|---|
| main areas (R) | hemi | subregions | | | | accuracy (mean) | cluster id |
| | | dk region | vertices | % of cluster | % of region | | |
| Temporo-parietal | Right | G_pariet_inf-Supramar | 196 | 79.0323 | 19.939 | 0.505 | 1 |
| | Right | G_temp_sup-Plan_tempo | 22 | 8.87097 | 8.08824 | 0.505 | 1 |
| | Right | G_pariet_inf-Angular | 16 | 6.45161 | 1.67189 | 0.505 | 1 |
| | Right | S_interm_prim-Jensen | 14 | 5.64516 | 7.1066 | 0.505 | 1 |
| Temporal | Right | G_temporal_middle | 288 | 75.5906 | 37.8947 | 0.506 | 2 |
| | Right | S_temporal_inf | 61 | 16.0105 | 19.4268 | 0.506 | 2 |
| | Right | S_temporal_sup | 26 | 6.82415 | 1.35699 | 0.506 | 2 |
| Temporal | Left | G_temporal_middle | 194 | 28.7407 | 27.9137 | 0.506 | 3 |
| | Left | S_temporal_sup | 156 | 23.1111 | 9.25267 | 0.506 | 3 |
| | Left | G_temporal_inf | 153 | 22.6667 | 23.6111 | 0.506 | 3 |
| | Left | S_oc-temp_lat | 93 | 13.7778 | 40.2597 | 0.506 | 3 |
| | Left | S_temporal_inf | 76 | 11.2593 | 23.1707 | 0.506 | 3 |
| Parietal | Left | G_pariet_inf-Supramar | 240 | 91.954 | 21.2578 | 0.506 | 4 |
| Orbitofrontal / frontopolar | Right | G_front_inf-Orbital | 74 | 33.945 | 84.0909 | 0.505 | 5 |
| | Right | Lat_Fis-ant-Horizont | 51 | 23.3945 | 41.8033 | 0.505 | 5 |
| | Right | G_front_inf-Triangul | 37 | 16.9725 | 13.4545 | 0.505 | 5 |
| | Right | G_orbital | 30 | 13.7615 | 4.47761 | 0.505 | 5 |
| | Right | S_orbital_lateral | 26 | 11.9266 | 26.8041 | 0.505 | 5 |
| Posterior insula | Right | Lat_Fis-post | 280 | 60.7375 | 46.9012 | 0.506 | 6 |
| | Right | S_circular_insula_inf | 101 | 21.9089 | 20.5703 | 0.506 | 6 |
| | Right | G_temp_sup-G_T_transv | 66 | 14.3167 | 42.8571 | 0.506 | 6 |
| Orbitofrontal / frontopolar | Left | S_orbital-H_Shaped | 109 | 22.7557 | 36.0927 | 0.505 | 7 |
| | Left | Lat_Fis-ant-Horizont | 101 | 21.0856 | 93.5185 | 0.505 | 7 |
| | Left | G_front_inf-Triangul | 73 | 15.2401 | 27.9693 | 0.505 | 7 |
| | Left | G_front_inf-Orbital | 65 | 13.5699 | 55.5556 | 0.505 | 7 |
| | Left | G_orbital | 65 | 13.5699 | 10.5863 | 0.505 | 7 |
| | Left | S_circular_insula_sup | 62 | 12.9436 | 8.21192 | 0.505 | 7 |
| Cingulate / medial frontal | Left | G_front_sup | 507 | 63.7736 | 25.0866 | 0.506 | 8 |
| | Left | G_and_S_cingul-Ant | 155 | 19.4969 | 23.7366 | 0.506 | 8 |
| | Left | G_and_S_transv_frontopol | 67 | 8.42767 | 46.2069 | 0.506 | 8 |
| | Left | G_front_middle | 44 | 5.53459 | 4.20248 | 0.506 | 8 |
| Superior frontal | Left | S_front_sup | 243 | 56.7757 | 30.8767 | 0.505 | 9 |
| | Left | G_front_sup | 104 | 24.2991 | 5.14597 | 0.505 | 9 |
| | Left | G_front_middle | 81 | 18.9252 | 7.73639 | 0.505 | 9 |

**Table S4: Cluster listings for taste category decoding.** For each cluster from the encoding, retrieval and cross-phase taste category searchlight analysis (main Fig. 5); cluster ID, mean decoding accuracy (chance=1/2), hemisphere, and the anatomical regions it spans. Regions are given as Destrieux atlas parcels (aparc.a2009s), grouped under broader anatomical headings for readability. For each parcel, the percentage of the cluster those vertices represent, and the percentage of the parcel falling within the cluster. Parcels are listed if they account for at least 5% of the cluster or if at least 50% of the parcel falls within the cluster.

| Encoding | | | | | | | |
|---|---|---|---|---|---|---|---|
| main areas (R) | hemi | subregions | | | | accuracy (mean) | cluster id |
| | | dk region | vertices | % of cluster | % of region | | |
| Lateral frontal | right | S_central | 599 | 7.7 | 52.0 | 0.56 | 1 |
| | right | G_front_sup | 1051 | 13.5 | 53.4 | 0.56 | 1 |
| | right | G_precentral | 653 | 8.4 | 70.4 | 0.56 | 1 |
| | right | G_front_middle | 551 | 7.1 | 54.3 | 0.56 | 1 |
| | right | S_front_middle | 501 | 6.4 | 98.4 | 0.56 | 1 |
| | right | G_front_inf-Triangul | 232 | 3.0 | 84.4 | 0.56 | 1 |
| | right | G_and_S_frontomargin | 213 | 2.7 | 97.7 | 0.56 | 1 |
| Cingulate | right | G_and_S_cingul-Mid-Ant | 284 | 3.6 | 52.1 | 0.56 | 1 |
| Orbitofrontal / frontopolar | right | S_orbital-H_Shaped | 224 | 2.9 | 64.2 | 0.56 | 1 |
| | right | S_orbital_lateral | 97 | 1.2 | 100.0 | 0.56 | 1 |
| | right | G_front_inf-Orbital | 79 | 1.0 | 89.8 | 0.56 | 1 |
| | right | G_and_S_transv_frontopol | 219 | 2.8 | 100.0 | 0.56 | 1 |
| Operculo-insular | right | Lat_Fis-ant-Vertical | 65 | 0.8 | 91.5 | 0.56 | 1 |
| | right | Lat_Fis-ant-Horizont | 106 | 1.4 | 86.9 | 0.56 | 1 |
| | right | G_front_inf-Opercular | 207 | 2.7 | 58.0 | 0.56 | 1 |
| Lateral frontal | Left | G_front_sup | 1547 | 8.9 | 76.5 | 0.556 | 2 |
| | Left | S_central | 787 | 4.5 | 65.7 | 0.556 | 2 |
| | Left | G_postcentral | 713 | 4.1 | 79.6 | 0.556 | 2 |
| | Left | G_precentral | 644 | 3.7 | 65.1 | 0.556 | 2 |
| | Left | G_and_S_subcentral | 565 | 3.3 | 97.8 | 0.556 | 2 |
| | Left | G_front_middle | 552 | 3.2 | 52.7 | 0.556 | 2 |
| | Left | S_front_inf | 498 | 2.9 | 88.0 | 0.556 | 2 |
| | Left | S_precentral-inf-part | 371 | 2.1 | 76.0 | 0.556 | 2 |
| | Left | G_and_S_paracentral | 335 | 1.9 | 53.1 | 0.556 | 2 |
| | Left | G_and_S_frontomargin | 234 | 1.4 | 100.0 | 0.556 | 2 |
| | Left | G_front_inf-Triangul | 229 | 1.3 | 87.7 | 0.556 | 2 |
| | Left | S_front_middle | 217 | 1.3 | 58.2 | 0.556 | 2 |
| Cingulate | Left | G_and_S_cingul-Mid-Post | 397 | 2.3 | 65.3 | 0.556 | 2 |
| | Left | G_and_S_cingul-Ant | 382 | 2.2 | 58.5 | 0.556 | 2 |
| | Left | G_and_S_cingul-Mid-Ant | 278 | 1.6 | 55.8 | 0.556 | 2 |

| | | | | | | | |
|---|---|---|---|---|---|---|---|
| Orbitofrontal / frontopolar | Left | S_orbital-H_Shaped | 178 | 1.0 | 58.9 | 0.556 | 2 |
| | Left | G_and_S_transv_frontopol | 145 | 0.8 | 100.0 | 0.556 | 2 |
| | Left | G_rectus | 156 | 0.9 | 53.4 | 0.556 | 2 |
| | Left | S_orbital_lateral | 78 | 0.5 | 100.0 | 0.556 | 2 |
| Operculo-insular | Left | G_Ins_lg_and_S_cent_ins | 124 | 0.7 | 59.3 | 0.556 | 2 |
| | Left | G_insular_short | 115 | 0.7 | 50.9 | 0.556 | 2 |
| | Left | G_front_inf-Opercular | 431 | 2.5 | 92.5 | 0.556 | 2 |
| | Left | Lat_Fis-ant-Vertical | 84 | 0.5 | 88.4 | 0.556 | 2 |
| | Left | S_circular_insula_sup | 659 | 3.8 | 87.3 | 0.556 | 2 |
| Temporal | Left | S_temporal_sup | 963 | 5.6 | 57.1 | 0.556 | 2 |
| | Left | G_temp_sup-Plan_tempo | 223 | 1.3 | 64.6 | 0.556 | 2 |
| | Left | G_temp_sup-G_T_transv | 177 | 1.0 | 91.7 | 0.556 | 2 |
| | Left | G_temp_sup-Plan_polar | 128 | 0.7 | 59.5 | 0.556 | 2 |
| | Left | S_temporal_transverse | 109 | 0.6 | 89.3 | 0.556 | 2 |
| | Left | G_temporal_middle | 400 | 2.3 | 57.6 | 0.556 | 2 |
| | Left | Pole_temporal | 390 | 2.3 | 94.0 | 0.556 | 2 |
| | Left | G_temp_sup-Lateral | 574 | 3.3 | 85.9 | 0.556 | 2 |
| Parietal | Left | G_parietal_sup | 665 | 3.8 | 66.1 | 0.556 | 2 |
| | Left | G_pariet_inf-Supramar | 568 | 3.3 | 50.3 | 0.556 | 2 |

| **Retrieval** | | | | | | | |
|---|---|---|---|---|---|---|---|
| **main areas (R)** | **hemi** | **subregions** | | | | **accuracy (mean)** | **cluster id** |
| | | **dk region** | **vertices** | **% of cluster** | **% of region** | | |
| Temporo-parietal | Right | S_temporal_sup | 377 | 68.8 | 19.7 | 0.52 | 1 |
| | Right | G_pariet_inf-Angular | 91 | 16.6 | 9.5 | 0.52 | 1 |
| | Right | S_interm_prim-Jensen | 41 | 7.5 | 20.8 | 0.52 | 1 |
| | Right | G_temp_sup-Lateral | 39 | 7.1 | 6.3 | 0.52 | 1 |
| Temporal pole | Right | Pole_temporal | 228 | 100.0 | 47.0 | 0.52 | 2 |
| Temporal | Left | S_oc-temp_med_and_Lingual | 215 | 51.9 | 33.1 | 0.524 | 3 |
| | Left | G_oc-temp_lat-fusifor | 196 | 47.3 | 44.4 | 0.524 | 3 |
| Parieto-occipital | Right | S_oc_sup_and_transversal | 198 | 39.8 | 52.2 | 0.522 | 4 |
| | Right | G_pariet_inf-Angular | 194 | 39.0 | 20.3 | 0.522 | 4 |
| | Right | G_occipital_middle | 69 | 13.9 | 13.2 | 0.522 | 4 |
| | Right | S_intrapariet_and_P_trans | 37 | 7.4 | 3.2 | 0.522 | 4 |
| Peri-central | Right | G_precentral | 293 | 74.7 | 31.6 | 0.52 | 5 |
| | Right | S_central | 99 | 25.3 | 8.6 | 0.52 | 5 |
| Operculo-insular | Right | S_circular_insula_sup | 274 | 79.4 | 44.7 | 0.522 | 6 |
| | Right | G_front_inf-Opercular | 24 | 7.0 | 6.7 | 0.522 | 6 |
| | Right | Lat_Fis-ant-Vertical | 20 | 5.8 | 28.2 | 0.522 | 6 |
| Cingulate / medial frontal | Left | G_front_sup | 971 | 59.6 | 48.0 | 522 | 7 |
| | Left | G_and_S_cingul-Mid-Ant | 292 | 17.9 | 58.6 | 522 | 7 |
| | Left | G_and_S_paracentral | 170 | 10.4 | 26.9 | 522 | 7 |
| | Left | S_cingul-Marginalis | 135 | 8.3 | 27.6 | 522 | 7 |

| Cross-phase | | | | | | | |
|---|---|---|---|---|---|---|---|
| main areas (R) | hemi | subregions | | | | accuracy (mean) | cluster id |
| | | dk region | vertices | % of cluster | % of region | | |
| Superior frontal | Right | G_front_sup | 256 | 45.4 | 13.0 | 0.516 | 1 |
| | Right | G_and_S_cingul-Ant | 142 | 25.2 | 18.8 | 0.516 | 1 |
| | Right | G_and_S_cingul-Mid-Ant | 120 | 21.3 | 22.0 | 0.516 | 1 |
| Anterior frontal | Right | G_front_sup | 95 | 30.4 | 4.8 | 0.513 | 2 |
| | Right | G_and_S_transv_frontopol | 58 | 18.6 | 26.5 | 0.513 | 2 |
| | Right | G_and_S_cingul-Ant | 48 | 15.4 | 6.3 | 0.513 | 2 |
| | Right | G_rectus | 37 | 11.9 | 15.7 | 0.513 | 2 |
| | Right | G_and_S_frontomargin | 35 | 11.2 | 16.1 | 0.513 | 2 |
| | Right | S_suborbital | 35 | 11.2 | 68.6 | 0.513 | 2 |
| Lateral prefrontal | Left | G_front_middle | 166 | 48.3 | 15.9 | 0.515 | 3 |
| | Left | S_front_inf | 138 | 40.1 | 24.4 | 0.515 | 3 |
| | Left | G_front_inf-Opercular | 27 | 7.8 | 5.8 | 0.515 | 3 |
| Temporal | Right | G_temporal_middle | 217 | 53.7 | 28.6 | 0.52 | 4 |
| | Right | S_temporal_sup | 121 | 30.0 | 6.3 | 0.52 | 4 |
| | Right | S_temporal_inf | 49 | 12.1 | 15.6 | 0.52 | 4 |
| Temporal | Left | G_temp_sup-Lateral | 142 | 29.0 | 21.3 | 0.52 | 5 |
| | Left | G_temporal_middle | 124 | 25.4 | 17.8 | 0.52 | 5 |
| | Left | S_temporal_sup | 94 | 19.2 | 5.6 | 0.52 | 5 |
| | Left | G_temp_sup-Plan_polar | 90 | 18.4 | 41.9 | 0.52 | 5 |
| Posterior cingulate | Right | S_cingul-Marginalis | 164 | 43.3 | 27.4 | 0.517 | 6 |
| | Right | S_subparietal | 116 | 30.6 | 22.8 | 0.517 | 6 |
| | Right | G_cingul-Post-dorsal | 82 | 21.6 | 39.4 | 0.517 | 6 |
| Posterior cingulate | Left | G_cingul-Post-dorsal | 159 | 25.9 | 63.9 | 0.515 | 7 |
| | Left | G_precuneus | 111 | 18.1 | 12.4 | 0.515 | 7 |
| | Left | S_pericallosal | 110 | 17.9 | 20.8 | 0.515 | 7 |
| | Left | G_cingul-Post-ventral | 96 | 15.7 | 68.1 | 0.515 | 7 |
| | Left | S_parieto_occipital | 76 | 12.4 | 14.0 | 0.515 | 7 |
| | Left | S_subparietal | 60 | 9.8 | 16.9 | 0.515 | 7 |
| Parieto-occiptal | Right | G_cuneus | 167 | 30.3 | 43.6 | 0.517 | 8 |
| | Right | S_parieto_occipital | 160 | 29.0 | 31.1 | 0.517 | 8 |
| | Right | G_occipital_sup | 115 | 20.8 | 29.3 | 0.517 | 8 |
| | Right | G_precuneus | 80 | 14.5 | 8.2 | 0.517 | 8 |
| | Right | G_parietal_sup | 30 | 5.4 | 3.8 | 0.517 | 8 |
| Temporo-parietal | Right | G_pariet_inf-Angular | 313 | 62.7 | 32.7 | 0.516 | 9 |
| | Right | S_intrapariet_and_P_trans | 183 | 36.7 | 16.0 | 0.516 | 9 |
| Temporo-parietal | Left | G_pariet_inf-Supramar | 365 | 79.5 | 32.3 | 0.516 | 10 |
| | Left | S_interm_prim-Jensen | 38 | 8.3 | 29.0 | 0.516 | 10 |
| | Left | G_temp_sup-Plan_tempo | 37 | 8.1 | 10.7 | 0.516 | 10 |
